# *In vivo* transcriptome analysis provides insights into host-dependent expression of virulence factors by *Yersinia entomophaga* MH96, during infection of *Galleria mellonella*

**DOI:** 10.1101/2020.08.31.276279

**Authors:** Amber R. Paulson, Maureen O’Callaghan, Xue-Xian Zhang, Paul B. Rainey, Mark R.H. Hurst

## Abstract

The function of microbes can be inferred from knowledge of genes specifically expressed in natural environments. Here we report the *in vivo* transcriptome of the entomopathogenic bacterium *Yersinia entomophaga* MH96, captured during three stages of intrahemocoelic infection in *Galleria mellonella*. A total of 1,285 genes were significantly upregulated by MH96 during infection; 829 genes responded to *in vivo* conditions during at least one stage of infection, 289 responded during two stages of infection and 167 transcripts responded throughout all three stages of infection. Genes upregulated during the earliest infection stage included components of the insecticidal toxin complex Yen-TC (*chi1, chi2* and *yenC1*), genes for rearrangement hotspot element containing protein *yenC3*, cytolethal distending toxin *cdtAB* and vegetative insecticidal toxin *vip2*. Genes more highly expressed throughout the infection cycle included the putative heat-stable enterotoxin *yenT* and three adhesins (usher-chaperone fimbria, filamentous hemagglutinin and an AidA-like secreted adhesin). Clustering and functional enrichment of gene expression data also revealed expression of genes encoding type III and VI secretion system-associated effectors. Together these data provide insight into the pathobiology of MH96 and serve as an important resource supporting efforts to identify novel insecticidal agents.

## Introduction

For successful invasion, colonization and bioconversion of host tissues, bacterial pathogens must acclimate to the host environment via alterations in patterns of gene expression (Crawford et al., 2010; Fang et al., 2016; Heroven & Dersch, 2014; Reniere, 2018; Tran et al., 2013). Knowledge of those genes activated specifically within host tissues allows insight into the genetic determinants of *in vivo* colonization and pathogenicity. An often used strategy involves transcriptional analysis of pathogen gene expression (transcriptomics) based on patterns of mRNA transcript abundance during infection (Arenas et al., 2019; Deng et al., 2018; Fan et al., 2020; Griesenauer et al., 2019; Haueisen et al., 2019; Ibberson & Whiteley, 2019; Luo et al., 2020; Nobori et al., 2018; Sun et al., 2019; Tang et al., 2019; Valenzuela-Miranda & Gallardo-Escárate, 2018).

*In vivo* transcriptomic approaches have been widely applied to human pathogenic bacteria using cell lines and model hosts (Avican et al., 2015; Baddal et al., 2016; Damron et al., 2016; Nuss et al., 2017; Westermann et al., 2016; Yan et al., 2013). Additionally, time-resolved *in vivo* transcriptomics have increased understanding of the function of bacterial pathogens during natural infection processes (Aprianto et al., 2016; Li et al., 2020; Luo et al., 2020; Tang et al., 2020; Westermann et al., 2019). However, similar strategies are yet to be applied to pathogens infecting insects, despite the obvious potential to identify new insect-active agents.

*Yersinia entomophaga* MH96 is an entomopathogenic bacterium that was isolated from *Costelytra giveni* (Coleoptera: Scarabaeidae) (Hurst et al., 2011), an endemic and economically significant pasture pest in New Zealand (Ferguson et al., 2019). Development of MH96 as a biopesticide has proven consistent pathogenesis by *per os* challenge against *C. giveni*, as well as a wide range of coleopteran, lepidopteran and orthopteran species (Hurst et al., 2015, 2011, 2019, 2014; Marshall et al., 2012). The key virulence determinant of MH96 is an insecticidal toxin complex (TC) called Yen-TC, which facilitates ingress of MH96 through the insect midgut (Hurst et al., 2011, 2014; Marshall et al., 2012) and is constitutively produced and secreted by MH96 when grown in broth culture at temperatures of 25 °C or lower (Hurst et al., 2011). In addition to the orally active insecticidal activities of Yen-TC, structural characterization and understanding the molecular mechanisms of pore-formation and translocation of cytotoxin has been the focus of much research (Busby et al., 2012, 2013; Landsberg et al., 2011; Piper et al., 2019).

Despite the importance of Yen-TC in MH96’s ability to breach the insect-midgut, additional investigations have suggested that as yet unknown virulence factors (VFs) are deployed during intrahemocoelic infection of *G. mellonella*. As a prime example, Hurst et al. 2015 showed that both wild-type MH96 and ΔTC (a Yen-TC deficient strain) have the same median lethal dose (LD_50_) of only ∼ 3 cells and were equally as effective at stopping the host’s phenoloxidase immune response when injected directly into the hemocoel of *G. mellonella* (Hurst et al., 2015). Genes for many other diverse putative VFs, including known insecticidal toxins and effectors, secretion and iron acquisition systems, proteolytic enzymes and adhesins have also been identified in the MH96 draft genome (Hurst et al., 2016). The extensive repertoire of putative VF-encoding genes represents an opportunity to utilize *in vivo* transcriptomics to further characterize this entomopathogen’s mode of action against insects, especially during intrahemocoelic infection of *G. mellonella*.

Larvae of the greater wax moth *G. mellonella* (Lepidoptera) have emerged as a valuable model host for study of microbial infection and innate immunity (Champion et al., 2016; Desbois & McMillan, 2015; Tsai et al., 2016). Yet, to date, pathogen *in vivo* transcriptome studies from *G. mellonella* have been confined to fungal pathogens only, including *Candida albicans* (Amorim-Vaz et al., 2015) and entomopathogenic *Beauveria bassiana* (Chen et al., 2018). Since the immune system of *G. mellonella* consists of both cellular (i.e., hemocytes) and humoral components (Singkum et al., 2019; Wojda, 2017), the intrahemocoelic infection is useful to characterize host-dependent genome-wide transcriptional modulation by MH96 in response to three stages of infection using a time-resolved approach.

Here, we focus on the interaction between MH96 and *G. mellonella*, using an *in vivo* transcriptomic strategy to identify pathogen genes activated under insect immunity, septicemia and pre-cadaver stages of intrahemocoelic infection. Differential expression (DE) analysis comparing MH96 transcriptome under *in vivo* and *in vitro* conditions identified pathogen genes, including known and putative VFs, that significantly responded either at specific or multiple timepoints throughout the infection process. To further understand the pathobiology of MH96, time-resolved *in vivo* versus *in vitro* fold-changes were clustered and functionally annotated using a database of known bacterial VFs. Particularly notable was upregulation of multiple genes encoding predicted adhesins, toxins and type III secretion system (T3SS) and T6SS translocated effectors.

## Materials and Methods

### *Intrahemocoelic infection of* G. mellonella *and pathogen RNA enrichmen*

Larval *G. mellonella* (Biosuppliers, Auckland, New Zealand) were provided with fresh diet of liquid honey, glycerol, Farex® brand baby rice cereal and granular yeast for up to one week. Healthy final instar larvae of similar lengths (> 20 mm, between 0.15 – 0.30 g) were selected and immobilized on ice. For intrahemocoelic infection, MH96 was grown overnight (18 h) in 3 ml Luria Broth Base (Invitrogen) with shaking (250 rpm) at 30 °C and then cultures were diluted in phosphate buffer solution (PBS, Sigma) and placed on ice prior to injection. Larvae were surface sterilized with 70 % ethanol and then injected below the third right leg with 10 μl of inoculum using a 30 gauge 12 mm needle on a 1 ml tuberculin syringe (Terumo) with a micro-injector.

Dilutions of MH96 ranging from ∼ 157 up to 2 × 10^7^ colony forming units (CFUs) were injected into *G. mellonella* depending on the stage of infection for each *in vivo* treatment (i.e., early, middle or late) (supplemental Table S1). For the early infection treatment, *G. mellonella* were injected with approximately 2 × 10^7^ CFUs and hemolymph was collected within 75 – 90 min post-infection, which was considered sufficient time to trigger a measurable host-dependent transcriptional response by MH96 under insect immunity. With respect to middle and late *in vivo* treatments, a 48 hours-post infection (HPI) growth curve was first obtained. MH96 CFU/larva were enumerated by dilution plating of *G. mellonella* homogenate on *Yersinia* semi-selective medium at specific time-points following injection of ∼ 345 CFUs (Supplementary Figure S1). This enabled definition of the 18-19 and 26-30 h incubation times for middle (i.e., septicemia) and late infection (i.e. pre-cadaver) treatments, respectively. Also, similar to the 48 HPI growth curve, lower inocula dosages of ∼157 and 693 CFUs were injected for the middle and late infection treatments, respectively compared to the early infection.

Following injection, the larvae were placed in petri-dishes and incubated at 25 °C within sealed plastic bags with moistened paper towel. After incubation, hemolymph was obtained by puncturing the dorsal cuticle near the second segment with a needle. Hemolymph was pooled from five to six individuals, yielding approximately 150 μl total volume. Four biological replicate samples were collected from each of the three *in vivo* infection stages (12 *in vivo* samples total). Pooled hemolymph samples were collected directly into microcentrifuge tubes with 300 μl of RNAprotect Bacterial Reagent (Qiagen), vortexed for five seconds, incubated at room temperature for 5 min and then centrifuged at 300 x g for 5 min at 4 °C to pellet hemocytes and cell debris. The supernatant was carefully separated from the host cell pellet by pipette and bacterial cells were then pelleted by centrifugation (10 min at 5,000 x g at ambient temperature (∼ 22 °C)) and the supernatant removed and stored at -80 °C. To estimate the CFUs per larva, serial dilutions from *G. mellonella* homogenate were plated on a *Yersinia-*semi selective medium and all *Yersinia-*positive colonies were counted within 24 h (5 separate larva per treatment; surface sterilized with 70 % ethanol). The weight of each larva was measured enabling determination of CFU per gram larval homogenate.

Control *in vitro* RNA samples were collected separately from broth cultures and two independent biological replicate samples were collected for each of the *in vitro* treatments – lag, exponential and stationary growth phases (6 *in vitro* samples total). For *in vitro* growth MH96 was grown overnight in 3 ml broth at 30 °C with shaking (250 rpm). The next day, flasks containing 50 ml fresh broth were inoculated with 1 % overnight culture and grown at 25 °C until samples were collected after 4 h 45 min (2 ml), 6 h (1 ml) and 9 h (1 ml) corresponding to specific cell densities of 3.2 ± 0.1 × 10^7^, 2.9 ± 0.9 × 10^8^ and 4.4 ± 0.1 × 10^9^ CFU ml^-1^ (comparable to CFU g^-1^ assessed from *in vivo G. mellonella* hemolymph RNA-seq samples), respectively. *In vitro* RNA samples were stabilized in RNAprotect Bacterial Reagent (Qiagen) according to the manufacturer’s protocol.

### RNA Extraction, library preparation and sequencing

Within one week of collection, RNA samples were re-suspended in 100 μL Tris-EDTA buffer (10mM Tris-HCl, 1mM EDTA; pH 8.0) (Ambion) containing 15 mg ml^-1^ lysozyme (Roche or Sigma-Aldrich) and 10 μl proteinase K (Roche). The samples were incubated for 10 min, interrupted by 10 s vortex every two min to lyse the cells. The RNeasy mini kit (Qiagen) was used for RNA extraction and RNA clean-up, which included two separate DNA digestions (on-column then off-column) with DNAse (Qiagen), all according to manufacturer’s guidelines. RNA samples were then precipitated at -20 °C overnight in 3 volumes 100 % cold isopropanol with 1/10 volumes 3M sodium acetate (pH 5.2), followed by two 70 % cold ethanol washes. Purified RNA was re-suspended in RNAse-free water and stored at -80 °C.

RNA extract from early infection hemolymph contained approximately half the amount of total RNA required for sequencing, so early infection RNA samples were doubled such that each sample was comprised from a pool of 10 - 12 larvae, while middle and late infection RNA samples were from five or six pooled individuals. Sufficient removal of genomic DNA from the samples was determined by confirming lack of PCR amplification from *in vivo* RNA template using primers RecA_352F 5’ – TCTCAGCCAGATACCGGTGA and RecA_987R 5’-CAGCAACATTTCACGCAGCT for the housekeeping gene *recA*. Middle and late infection *in vivo* RNA samples containing ∼7.2 μg total RNA and early infection samples containing < 7.2 μg were stabilized on RNAstable 1.5 micro-centrifuge tubes (Biomātrica) by SpeedVac (Savant) at ambient temperature (∼ 22 °C) for 1.5 h.

**Table 1:** MH96 cell density under *in vivo* and *in vitro* conditions used for differential expression analysis.

| *In vivo* infection stage/*in vitro* growth | Mean cell density *in vivo* (i.e., *G. mellonella* homogenate) (CFU g<sup>-1</sup> ± SEM<sup>b</sup>) | Mean cell density *in vitro* (i.e., Luria Broth Base broth) (CFU ml<sup>-1</sup> ± SEM) |
| --- | --- | --- |
| Early infection/lag-growth | $4.4 \pm 0.5 \times 10^7$ | $3.2 \pm 0.1 \times 10^7$ |
| Middle infection/exponential growth | $4.4 \pm 1.7 \times 10^8$ | $2.9 \pm 0.9 \times 10^8$ |
| Late infection/stationary growth | $5.0 \pm 2.2 \times 10^9$ | $4.4 \pm 0.1 \times 10^9$ |

<sup>a</sup>CFU = colony forming units; <sup>b</sup>SEM = standard error of the mean.

Macrogen Korea carried out library preparation and sequencing. Following recovery from the RNAstable tubes, RNA quantity and Integrity Number were determined using the 2200 TapeStation (Agilent Technologies). Host and bacterial ribosomal RNA (rRNA) was depleted from the *in vivo* samples using the ScriptSeq Complete Gold Kit (Epidemiology) (Illumina) then strand-specific cDNA libraries were prepared using the ScriptSeq RNA-Seq Library Preparation Kit (Illumina) according to the manufacturer’s guidelines. The *in vitro* samples were prepared using the TruSeq stranded mRNA (Illumina) with the Ribo-Zero rRNA removal kit for Bacteria (Epicentre) according to standard protocol. To verify the size of the PCR enriched fragments, the template size distribution was assessed on the 2100 Bioanalyzer using a DNA 1000 chip (Agilent Technologies). The libraries were sequenced on the HiSeq2500 platform (Illumina) (Version HCS v2.2) to generate 101 bp paired-end reads.

### Bioinformatic analysis

Adapter and barcode contamination were filtered using *Flexbar* (v2.4) against Illumina adapter and barcode reference sequences (supplemental Table S2 and Table S3). Reads were trimmed in ‘ANY’ mode with minimum overlap of five, maximum uncalled base of one and minimum quality threshold of 20 (Dodt et al., 2012). The bbsplit.sh function of the *BBMap* package was used to quantify and remove contaminating rRNA sequences with minimum ratio of 0.5, minimum hit number of one and maximum insertion/deletion length of 500 bp (Bushnell, 2015). All libraries were screened against full-length 5S, 16S and 23S rRNA sequence from the MH96 genome (see accession information below). To remove host sequences, *in vivo* libraries were also mapped against the *G. mellonella* partial 18S and 28S rRNA sequence retrieved from GenBank (Accessions: <u>U65198.1</u>, <u>U65138.1</u>, <u>AF286298.1</u>, <u>AF423921.1</u> and <u>X89491.1</u>) and the *G. mellonella* immune cDNA library (JG394435 – JG406465) (Vogel et al., 2011).

Remaining paired-end reads were aligned to the MH96 genome (Accession: CP010029.1, <u>GI: 1034308998</u>) using *Rockhopper* (v2.03) in verbose mode. Alignment stringencies were increased from default values, such that allowable mismatch threshold was decreased from 15 to 10 % of read length and the minimum seed was increased from 33 % to 50 % of read length. Any residual count data aligning to 5S, 16S and 23S rRNA genes or any short (less than 50 bp) non-coding RNA or anti-sense RNA were not considered further in the analysis.

The early infection *in vivo* libraries were investigated whether the sequence depth was estimated to have adequately captured the majority of MH96 low-abundance transcripts using the *iNEXT* R package (nboot = 50) (v2.0.18) (Hsieh et al., 2016). The *RUVseq* R package (v1.16.0) was used to generate relative log expression (RLE) plots and conduct principle component analysis (PCA) (Risso et al., 2014). Upper-quartile normalized (Bullard et al., 2010) libraries were filtered for transcripts with greater than 5 aligned reads in at least two libraries.

Normalized expression data were then converted to log2-counts-per-million (Log_2_CPM) using the R package *limma* (v3.38.3) (Ritchie et al., 2015; Smyth, 2004) and the mean-variance relationship was modeled using the ‘voom’ method (Law et al., 2014). Genome-wide host-dependent responses by MH96 to both *in vitro* and *in vivo* conditions were visualized with *Circos* (v0.69) (Krzywinski et al., 2009) and putative genomic islands (GIs) were assigned using the webserver <u>IslandViewer 4</u> (Bertelli et al., 2017). To determine MH96 genes that significantly responded to *in vivo* conditions a DE analysis was undertaken by comparing average transcript abundance between *in vivo* and *in vitro* libraries sharing equivalent cell densities (i.e., early infection vs. lag growth in broth, middle infection vs. exponential growth in broth and late infection vs. stationary growth in broth). Genes with significantly DE were defined by multiple testing with false discovery rate of 0.05 using the Benjamini and Yekutieli multiple test correction method (Benjamini & Yekutieli, 2001) using *limma*.

Time-resolved average *in vivo* vs. *in vitro* log_2_CPM fold-change values for all genes with significantly DE identified between the *in vivo* and *in vitro* treatments (DE analysis described above) were clustered using the fuzzy c-means (FCM) algorithm (Futschik, Matthias & Carlisle, 2005) with the *MFuzz* R package (v2.42.0) (Kumar & Futschik, 2007). A heuristic qualitative assessment was used to identify both the optimal FCM parameter (i.e., ‘fuzzifier’) and number of clusters (Kumar & Futschik, 2007). The assessment considered both predicted operon structure (output from *Rockhopper*) and functional annotations assigned using homology based search (BLASTx, e-value cut-off = 1 × 10^−8^) against the <u>virulence factor database</u> (VFDB) (accessed July, 2018) (Chen et al., 2016).

MH96 host-specific factors were identified among clusters containing transcripts with comparably greater median log_2_CPM *in vivo* fold-change values (≥ ∼ 2) for at least one stage of growth during intrahemocoelic infection. Next, these host-specific factors were functionally characterized as putative VFs when significant sequence homology was identified against an already characterized VF found in the VFDB. Finally, functional categories of virulence were assigned to all of these putative VFs primarily based on the VFDB-derived annotation, in accordance to our definitions found in supplemental Table S4. When VFDB-derived annotations were not informative, additional information such as, top-scoring BLASTx hits from the GenBank non-redundant protein database, were used to assign a possible function or otherwise these hypotheticals were left unclassified.

### Data availability

Sequence data and both raw and normalized count tables were deposited in the GenBank Geo Submission Omnibus under accession: <u>GSE142509</u>. All genes with significantly DE detected between *in vivo* versus *in vitro* time-resolved contrasts (early infection vs. lag growth *in vitro*; middle infection vs. exponential growth *in vitro*; late infection vs. stationary growth *in vitro*) are provided in supplementary Table S5. All functionally annotated putative VFs identified from host-specific fuzzy clusters are presented in supplementary Table S6. Open-source R code for statistical analysis is available at: https://github.com/damselflywingz/In_vivo_infection_series_fuzzy_RNA-seq.

## Results

### Experimental infection and RNA sequencing

MH96 was enriched from hemolymph at cell densities of 4.4 × 10^7^, 4.4 × 10^8^ and 5.0 × 10^9^ CFU g^-1^ (Table 1). In order to distinguish host-specific gene induction by MH96 from response to general growth conditions, *in vitro* samples were collected across three growth phases at approximately equivalent cell densities to the *in vivo* samples (Table 1).

After quality trimming, adapter removal of host and rRNA sequences, the paired-end reads were aligned to the draft MH96 genome. Only an average of 9 % of reads from early-infection libraries could be aligned to the reference, which was lower by comparison to the average reads aligned from middle (58 %) and late infection (59 %), or *in vitro* libraries (84 -93 %) (supplemental Figure S2 and Table S7). In order to determine if transcripts expressed in lower abundances were adequately captured within the early infection libraries, these libraries were analyzed by rarefaction, which showed that only a slight improvement in total transcript detection if sequencing depth were increased (supplemental Figure S3). Based on this analysis, the *in vivo* early infection libraries were determined to have captured most MH96 transcripts produced across a dynamic range of conditions expected under these conditions and were considered saturated.

Preliminary data analysis explored the libraries for unwanted technical variation arising from batch effects and potential outliers using both RLE plots and PCA (supplementary Figure S4 and Figure S5), respectively. Variation in sequencing depth between libraries was found to have affected the distribution of RLE values (supplementary Figure S4A), which was largely controlled for by upper quartile normalization (supplementary Figure S4B). Even after normalization, the variation in RLE values from the stationary growth *in vitro* libraries was greater when compared to all other libraries indicating residual unwanted technical variation arose from this treatment. A clear separation between *in vitro* and *in vivo* libraries was determined by PCA of normalized count data (supplemental Figure S5). Libraries consistently clustered by treatment, except one *in vivo* late infection library, which clustered more closely to the middle infection *in vivo* libraries and was identified as a biological outlier and removed from subsequent analysis.

### Differential expression of MH96 during early, middle and late infection stages

Comparison of average transcript abundances across three stages of growth identified 2,397 differentially expressed genes that were either up or downregulated between *in vivo* and *in vitro* libraries (supplemental Figure S6 and Table S5). MH96 upregulated 1,285 genes during intrahemocoelic infection of *G. mellonella*; 829 genes responded to *in vivo* conditions during at least one stage of infection, 289 responded during two stages of infection and 167 transcripts responded throughout all three stages of infection. MH96 also upregulated 1,243 genes under *in vitro* conditions, of which 131 were also found to be upregulated under *in vivo* conditions at different cell densities.

Regions of interest from the MH96 draft genome either known or suspected to encode VFs (supplemental Table S8) were explored with respect to time-resolved genome-wide host-dependent transcriptional responses during intrahemocoelic infection (Figure 1). Several predicted GIs partially or fully overlapped with previously identified genomic regions of interest (i.e., unique region 1 overlaps Is.2, Rhs2 overlaps Is.5, T3SSYE1 overlaps Is.6, Rhs5 overlaps Is.11, T3SSYE2 overlaps Is.12, Rhs4 overlaps Is.14, unique region 2 overlaps Is.16, T2SS overlaps Is.18 and Rhs3/T6SS overlaps Is.21).

**Figure 1:**
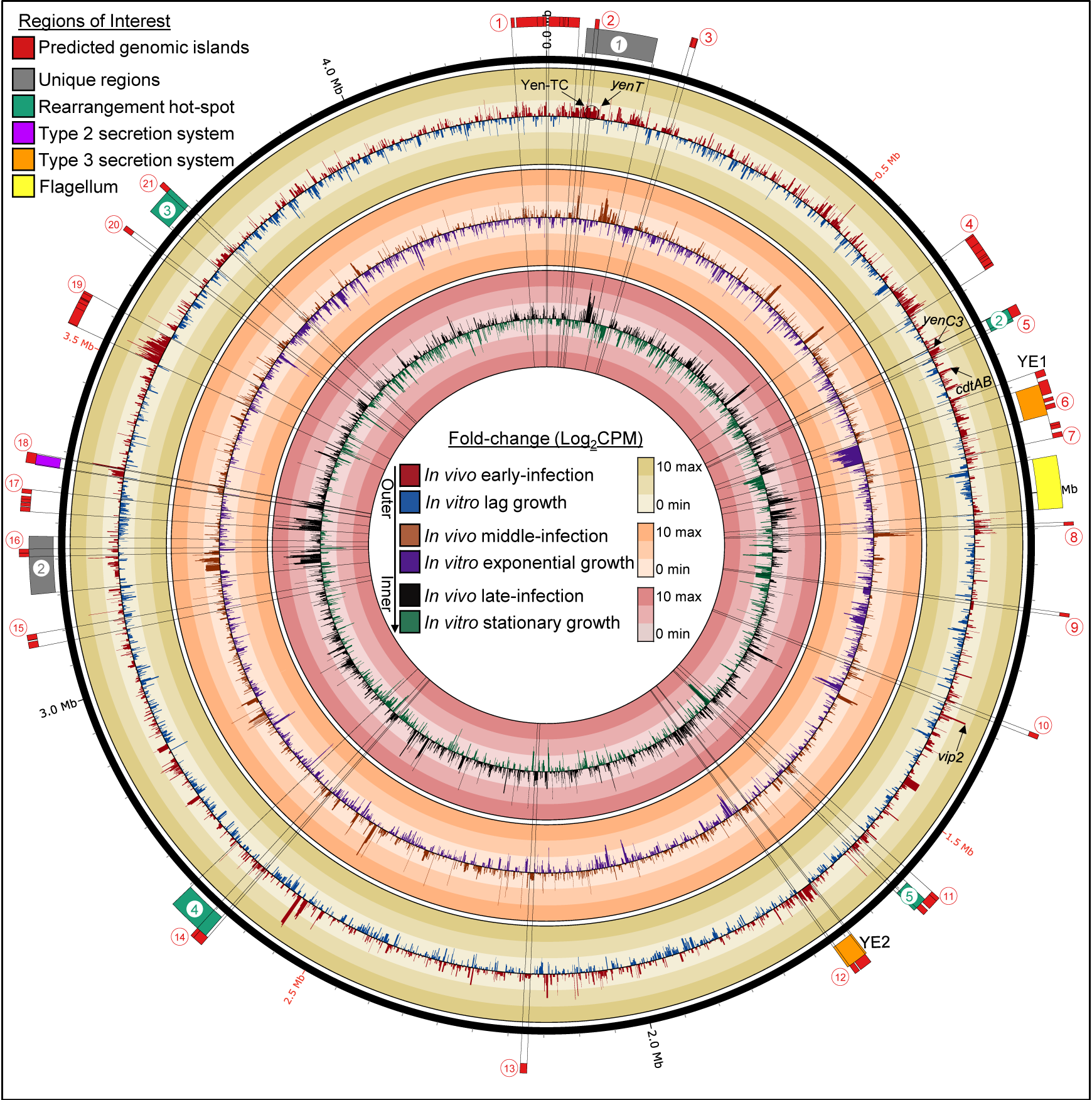
Circos plot of *Y. entomophaga* MH96 genome-wide transcriptional response to *in vivo* and *in vitro* conditions during early, middle and late infection and lag, exponential and stationary growth, respectively. CPM = Counts-per-million. Mean fold-change in log counts-per-million (Log_2_CPM) (*in vivo* early and middle infection: n = 4, *in vivo* late infection: n = 3 and all *in vitro*: n = 2). Red circles numbered 1 -21 represent predicted genomic islands identified using IslandViewer 4 webserver (Refer to supplemental Table S8). Regions of interest/loci previously identified from the draft genome are numbered or labelled as required (Hurst et al., 2016).

Wide-spread host-dependent upregulation of multi-gene clusters located within several regions of interest/predicted GIs, including unique regions 1 and 2, Rhs2/Is.5, T3SSYE2/Is.12 and T2SS/Is.18, was observed in MH96. Significantly higher expression for the entire T2SS island (PL78_08930 – PL78_08970) was measured during early and/or late infection stages compared to *in vitro* growth conditions. Significantly higher responses by genes for putative T3SSYE2 structural components (SpaS/PL78_14595, SpaP/PL78_14580, SpoA/PL78_14575, hypothetical protein/PL78_14570, SpaL/PL78_14560, InvA/PL78_14550, and PrgH/PL78_14530) to *in vivo* growth conditions compared to *in vitro* during early and/or late infection were also identified. In contrast, most of the T3SSYE1 structural component genes were significantly upregulated by MH96 either in response to *in vitro* conditions or both *in vitro* and *in vivo* conditions depending on cell density, except putative T3SSYE1-associated chaperone (PL78_18170) that was more highly expressed *in vivo* during early infection only. Genes located within a predicted phage-like island were also significantly upregulated during early stage of infection, compared to growth *in vitro* (Is.4: PL78_06170 – PL78_06360; PL78_19225 –PL78_19085, Is.8: PL78_17412 – PL78_17386 and Is. 19: PL78_06425 – PL78_06390; PL78_19127 – PL78_19160; PL78_00015 – PL78_00300).

Genes for other putative VFs from MH96 genomic regions of interest were more responsive during late stages of intrahemocoelic infection compared to other time points. Iron-acquisition gene clusters located within unique region 2 (PL78_09605, PL78_09620 – 09640, PL78_09665 – PL78_09710, PL78_09720, PL78_09730 – PL78_09745 and PL78_09780 – PL78_09785) were found to have higher expression under *in vivo* conditions, especially during later stages of infection. Genes for the flagellum (PL78_17795 – PL78_17785, PL78_17735 – PL78_17670, PL78_17655 – PL78_17610, PL78_17595 – PL78_17585) were also more highly expressed mostly during the later infection stage too. Further, phage-related and hypothetical genes encoded within a predicted GI Is.17 (PL78_09315 – PL78_0914) were also found to have significant transcriptional response to *in vivo* conditions compared to growth *in vitro* during late infection stages as well.

An important region in the MH96 genome is Unique Region 1 (contains PAI_YE96_) and genes for hypothetical proteins *yenU* (PL78_03775) and *yenW* (PL78_03790), putative heat-stable enterotoxin *yenT* (PL78_03785), iron-siderophore biosynthesis/transport (PL78_03905 – PL78_03955, excluding PL78_03935 and PL78_03940) and serralysin gene cluster (PL78_03960 – PL78_03995, excluding PL78_03985), were all more highly expressed in the host at various or throughout all stages of infection compared to *in vitro* treatments. Also located in Unique region 1, Yen-TC genes *chi1* (PL78_03740), *chi2* (PL78_03755) and *yenC1* (PL78_03765), were found to have significantly higher expression during early infection only, while DE was not detected for other Yen-TC component genes.

### Functional enrichment of *in vivo* host clusters

Qualitative assessment of fuzzy clustering results showed that 20 clusters with fuzzifier value of 1.5 was optimal for the analysis of fold-change log_2_CPM values comparing MH96 transcriptome across three different stages of intrahemocoelic infection to *in vitro* growth controls. Among these 20 clusters, eight clusters (containing 798 transcripts) exhibited the most striking responses to *in vivo* conditions compared to *in vitro* growth (supplemental Figure S7). These eight ‘*in vivo* clusters’ were further organized into three main ‘*in vivo* cluster types’ based on shared expression profiles across multiple *in vivo* clusters (Figure 2A). *In vivo* clusters containing transcripts with elevated abundances across all three stages of infection were identified as ‘Host All’ type clusters, while ‘Host Early’ and ‘Host Late’ type clusters were characterized by containing mostly transcripts with elevated abundances during early or late stages of infection only, respectively.

**Figure 2:**
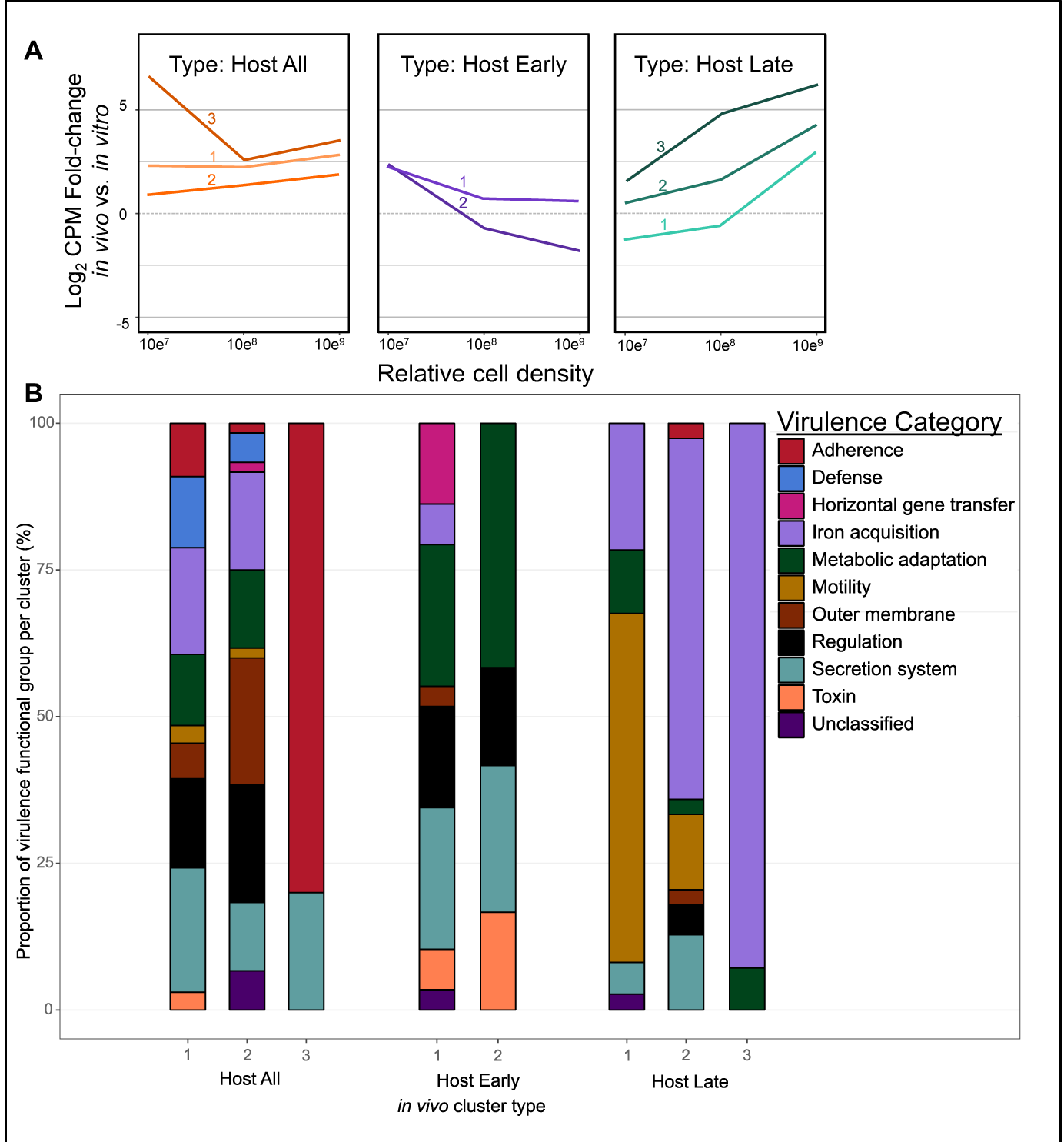
*Yersinia entomophaga* MH96 putative host-specific virulence factors (VFs) sharing significant sequence similarity to known VFs from the virulence factor database identified among host-specific *in vivo* clusters. A) Median log_2_ counts-per-million-fold-change between *in vivo* and *in vitro* libraries organized by *in vivo* cluster type (each line represents a separate host-specific *in vivo* cluster, refer to supplemental Figure S7). B) The proportion of genes organized by virulence function classification among *in vivo* clusters types. Refer to supplementary Table S6.

To further classify the role of putative MH96 VFs in pathogenesis, those transcripts from the eight *in vivo* clusters were functionally annotated using VFDB. Using this approach, 28.7 % of the transcripts from *in vivo* clusters could be further categorized by potential role in pathogenicity (i.e., adherence, defense response, horizontal gene transfer, iron acquisition, metabolic adaptation, mobility, outer membrane, regulation, secretion system and toxin/effector or unclassified) (Figure 2B, Figure 3 and supplemental Table S6). Transcripts related to iron acquisition, secretion systems, regulation and metabolic adaptation were commonly identified among each of the three different *in vivo* cluster types but an obvious temporal shift in the expression of iron acquisition and flagellar motility versus toxins, effectors and adhesions were identified when comparing early and late *in vivo* clusters.

**Figure 3:**
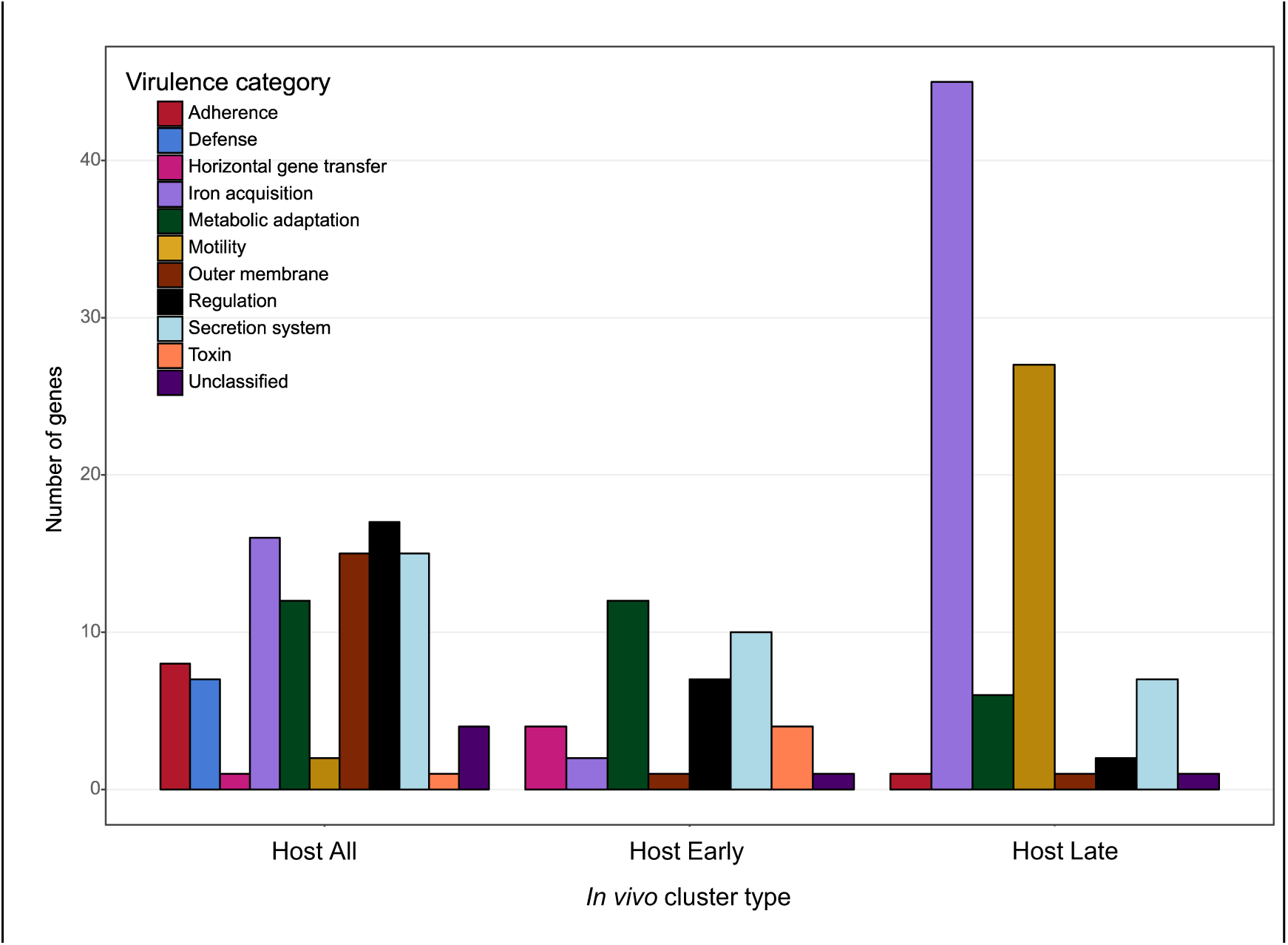
Number of genes assigned to each virulence category for putative virulence factors identified from *in vivo* cluster types. *In vivo* clusters combined by type: Host All = constitutively higher *in vivo* expression during earl-, middle and late infection; Host Early = higher *in vivo* during early infection only; Host Late = higher *in vivo* expression during late infection only. See Figure 2A for Host All, Host Early and Host Late median expression profiles and supplemental Table S6 for more details.

Four genes encoding an usher-chaperone fimbrial cluster (PL78_12465 – PL78_12480) co-expressed with putative T3SS lipoprotein chaperone *YscW-like* (PL78_14485), were identified as some of the most host-responsive genes in the MH96 genome, especially at the early stage of infection (*in vivo* early infection vs. *in vitro* growth fold-change change values ranging from greater than 4.6 up to 7.1x Log_2_CPM). Genes for filamentous hemagglutinin (PL78_11060, previously annotated as a hemolysin and its cognate two-partner secretion system (TPS) (PL78_11055) also had significantly higher expression during early and middle stages of intrahemocoelic infection. Also, a predicted fimbrial gene (PL78_08275) and chitin-binding protein (PL78_08295) both associated with the Chi-Fim region (Hurst et al., 2016) were also captured from *in vivo* Host all type clusters but the relative expression levels of the Chi-Fim-associated fimbrial and chitin-binding protein genes were orders of magnitude lower than genes for other putative MH96 adhesions that were upregulated *in vivo*.

Transcripts for several putative toxins and effectors were identified among *in vivo* clusters (Table 2). The aforementioned heat-stable enterotoxin *yenT* was the only putative toxin gene identified to be highly expressed *in vivo* at two timepoints during infection and the transcript encoding putative AidA-like type V secretion system (T5SS) secreted adhesin (PL78_10240) clustered with other MH96 transcripts with higher *in vivo* gene expression throughout infection (Table 2).

**Table 2:** Putative MH96 toxin and effector genes from host-specific gene expression clusters with significant sequence homology to other known toxins or effectors from VFDB

| Toxin/Effector type | Locus tag | Designation | Host cluster type | Region of interest/Genomic island/Locus <sup>a</sup> |
| --- | --- | --- | --- | --- |
| Toxin | PL78_03785 | *yenT* | Host all | Unique region 1-PAI<sub>YE96</sub> |
| Toxin | PL78_18780 | *yenC3* | Host early | Rhs-associated region 2/Is. 5 |
| Toxin | PL78_18440 | *cdtA* | Host early | *cdtAB* |
| Toxin | PL78_18445 | *cdtB* | Host early | *cdtAB* |
| Toxin | PL78_16145 | *vip2* | Host early | *vip2* |
| Effector - T3SS<sup>b</sup> | PL78_14610 | *sicC-like* | Host all | T3SSYE2/Is.12 |
| Effector - T3SS | PL78_07345 | *ipgD/sopB-like* | Host all | - |
| Effector - T3SS | PL78_14615 | *sicD-like* | Host early | T3SSYE2/Is.12 |
| Effector - T3SS | PL78_18085 | *ipaB/evcA-like* | Host early | T3SSYE1/Is.6 |
| Effector - T3SS | PL78_14295 | *sopD-like* | Host early | - |
| Tube/Effector - T6SS | PL78_02751 | *hpc* | Host all | - |
| Tube/Effector - T6SS | PL78_00900 | *hpc* | Host early | Rhs-associated region 3 |
| Effector - T6SS<sup>d</sup> | PL78_01040 | *rhs*-containing | Host early | Rhs-associated region 3/Is.21 |
| Tube/Effector - T6SS | PL78_03598 | *hpc* | Host late | Is.1 |
| Tube/Effector - T6SS | PL78_09211 | *hpc* | Host late | Is.17 |
| Effector-T5SS<sup>c</sup> | PL78_10240 | *aidA* | Host all | - |

<sup>a</sup>Regions of Interest (See Figure 1): MH96 pathogenicity-associated island (PAI<sub>YE96</sub>), rearrangement-hotspot (Rhs), MH96 type 3 secretion systems (T3SSs) T3SSYE1 and T3SSYE2 (Hurst et al., 2016); Predicted genomic islands see Figure 1 and supplementary Table S8; Loci: cyto-lethal distending toxin components (CdtA, CdtB), vegetative insecticidal protein 2 (Vip2) (Hurst et al., 2016); <sup>b</sup>T3SS = Type III secretion system; <sup>c</sup>T5SS = Type V secretion system and <sup>d</sup>T6SS = Type VI secretion system;

Several other transcripts for putative toxins were identified from Host Early *in vivo* clusters, including putative cytolethal distending toxin sub-components *cdtA* and *cdtB* (PL78_18440-PL78_18445), *yenC3* (PL78_18780, a YenC-homolog also known as *rhsA* encoded within Rhs2/Is.5) and putative vegetative insecticidal protein component, *vip2* (PL78_16145) (Table 2). Transcripts for known, and putative Yen-TC components, *chi1* and *yenC3* were captured within a Host Early *in vivo* cluster, respectively, but only *yenC3*, not *chi1*, shared any significant homology against VFs known found in the VFDB using BLASTx homology search.

In addition to predicted toxins, transcripts for T2SS and T3SSYE2 structural components were also identified as putative VFs from *in vivo* clusters (supplemental Table S6). Two genes for T3SSYE2 co-located putative effectors were represented from *in vivo* clusters and found to have significantly greater expression under *in vivo* conditions compared to *in vitro* controls, including SicC-like (PL78_14610) and SicD-like (PL78_14615), with the former being more highly expressed across all cell densities and the latter more highly expressed during early infection only. In addition to T3SSYE2-associated effector genes, two other genes for putative T3SS effectors were also found to be more highly expressed *in vivo* during early infection including T3SSYE1-associated IpaB/EvcA-like (PL78_18085) and orphan SopD-like (PL78_14295). Another gene for putative T3SS orphan effector, IpgD/SopB-like (PL78_07345) represents a fifth possible T3SS effector, with significantly increased expression throughout all three stages of infection, compared to *in vitro* controls.

With respect to T6SS, structural components for the machinery were not identified as having significantly greater expression *in vivo* compared to *in vitro* controls, but putative T6SS effector transcripts were identified from several of the *in vivo* cluster types, including Rhs-containing protein (PL78_01040) and five different hcp-type structural tube/effector proteins (PL78_02751, PL78_03598, PL78_03602, PL78_09211 and 00900). Also, the gene encoding the putative T6SS effector that shares sequence similarity to the eukaryotic transcription elongation factor domain Spt4 (PL78_00995) was found to have higher *in vivo* gene expression throughout infection but was not assigned a VFDB annotation based on amino acid sequence.

Not all MH96 predicted toxins and effectors were highly expressed under *in vivo* condition though; genes for putative PirAB binary toxin (PL78_09590 and PL78_09595), RTX toxin (PL78_16910) and adenylate cyclase (PL78_08395) were found to have higher expression during exponential and/or stationary growth *in vitro*, with no significant response to *in vivo* conditions observed. Furthermore, despite having significantly greater abundance during early-infection of *G. mellonella* compared to growth *in vitro*, transcripts for some other known or suspected MH96 VFs, including insecticidal TC components Chi2 and YenC1, and putative T3SS effector LopT (PL78_18760) were not found to cluster among the eight *in vivo* clusters, instead clustering with other transcripts associated with comparatively lower overall median *in vivo* vs. *in vitro* fold-change values.

## Discussion

Our *in vivo* transcriptome analysis has shed light on MH96 genes activated during intrahemocoelic infection of *G. mellonella*. Major shifts in gene expression by MH96 were observed between *in vivo* and *in vitro* conditions across all three stages of growth and the mosaic nature of the MH96 genome was highlighted by wide-spread co-expression of horizontally acquired genes for Yen-TC components and putative toxins (YenT, YenC3), translocated effectors (T3SS and T6SS), secretion systems (T2SS and T3SSYE2) and iron acquisition systems in response host-dependent conditions. These findings provide evidence that complex genome-wide responses by MH96 occur throughout different stages of insect infection and reveal important insights into host-dependent transcriptional responses by entomopathogenic bacteria in nature.

Upon entry to the hemocoel, entomopathogenic bacteria must respond to the cellular and humoral components of the insect host immune system (Nielsen-LeRoux et al., 2012). Within lepidopteran larvae, the hemolymph contains different hemocytes involved in phagocytosis, nodulation and encapsulation as well as humoral factors like prophenoloxidase, antimicrobial peptides, reactive nitrogen and oxidative species or lysozyme (Casanova-torres & Goodrich-blair, 2013; Jiang et al., 2010; Wojda, 2017). In order to occupy the insect hemocoel, MH96 produces VFs that combat the host immune defenses during early stages of infection, including anti-hemocytic factors such as toxins or translocated effectors. Upon incapacitation of the host immune system, MH96 produces additional VFs that function to mobilize and acquire essential nutrients, like iron, from insect cells or tissues to fuel pathogen metabolism and proliferation, ultimately leading to septicemia and host death (Nielsen-LeRoux et al., 2012).

During early stages of intrahemocoelic infection MH96 upregulates expression of Yen-TC components *chi1, chi2* and *yenC1* as well as putative Yen-TC component *yenC3* and toxins *cdtAB* and *vip2*, which are known or suspected to be anti-hemocytic factors based on functional enrichment. CdtAB is a predicted genotoxin, which are known to interfere with normal hemocyte cell cycle (Heywood et al., 2005; Lara-Tejero & Galán, 2000), while Vip2 and YenC3 may act on host cytoskeleton through (ADP)-ribosyltransferase activity targeting globular (G)-actin (Barth et al., 2004) or Rho GTPAses (Lang, Alexander et al., 2010), respectively. While Yen-TC is established as the primary orally-active virulence determinant of MH96 (Hurst et al., 2011; Marshall et al., 2012), purified Yen-TC was also found to cause ruffling of *G. mellonella* hemocytes (Hurst et al., 2015) and nuclear and actin condensation and apoptosis of *Spodoptera frugiperda* insect cell line Sf9 (Marshall et al., 2012). Taken together, Yen-TC should be considered both an anti-hemocytic factor as well as an orally-active insecticidal toxin.

Unlike putative toxin genes that were upregulated only during the early stage of infection, the MH96 heat stable enterotoxin, *yenT*, was highly expressed during both the middle and late stages of infection compared *in vitro* control treatments. Similar in size YenT shares *N*-terminal amino acid sequence homology to the heat-stable enterotoxins YstA and YstB (Hurst et al., 2011), encoded by gastrointestinal-disease causing strains of *Y. enterocolitica* (Delor et al., 1990; Grant et al., 1998; Ramamurthy et al., 1997; Singh & Virdi, 2004), *Y. frederiksenii, Y. intermedia* and *Y. kristensenii* (Imori et al., 2017). Recently the heat-stable enterotoxin, YacT, produced by *Y. frederiksenii* has been implicated in host immune modulation following intrahemocoelic injection of *G. mellonella*. The *E. coli* recombinant YacT, altered the morphology and increased the aggregation of *G. mellonella* hemocytes (Springer et al., 2018). It is plausible that, like YacT of *Y. frederiksenii*, MH96 YenT also functions as an anti-hemocytic factor given the responsiveness of *yenT* to intrahemocoelic conditions at multiple stages of infection.

Among pathogenic *Yersinia*, T3SSs are known to target host cells through tight attachment and translocate effectors into the cytoplasm to disrupt cytoskeleton, trigger apoptosis, prevent phagocytosis or interfere with immune pathways (Bliska et al., 2013; Pha & Navarro, 2016; Plano & Schesser, 2013). The *Y. enterocolitica* T3SS Ysa (found in biotype 1B strains) (Foultier et al., 2002; Thomson et al., 2006) shares the same gene order with MH96 T3SSYE2 (Hurst et al., 2016) of the predicted island Is.12, suggesting a common origin in both pathogens. In addition to interacting with mammalian cells (Bent et al., 2013; Matsumoto & Young, 2006), Ysa is essential for replication within *Drosophila melanogaster* S2 cells (Walker et al., 2013) and secretes effectors at temperatures of less than 26 °C in nutrient-rich broth with high salt concentrations (Venecia & Young, 2005). Based on the similar gene order to Ysa and the upregulation during intrahemocoelic infection, T3SSYE2 is likely another anti-hemocytic mechanism used by MH96 to overcome the insect immune system.

Also, similar to Ysa, which secretes effectors encoded by genes within distal regions of the *Y. enterocolitca* genome (Walker & Miller, 2009), the MH96 *in vivo* transcriptome revealed evidence for coordinated host-dependent regulation of genes for T3SSYE2 structural components and several predicted effectors, some of which were not co-located within the T3SSYE2 region of the genome. For example, MH96 encodes a gene for a putative LopT-like effector, which is homologous to LopT, a T3SS effector (similar to YopT of *Y. pestis*) with known insecticidal activity in *Photorhabdus luminescens* (Brugirard-Ricaud et al., 2005). Expression of the LopT-like effector gene was significantly upregulated by MH96 during early stage infection and notably, *lopT* and *yenC3* are both co-located within an effector-island, (i.e., Rhs-associated region 2/Is.5), and are possibly co-opted by respective translocation machineries T3SSYE2 and Yen-TC and reflect the multiple diverse modes of action deployed by MH96 against the host during early stages of intrahemocoelic infection.

In contrast to T3SSYE2 and its predicted effectors that were upregulated under *in vivo* conditions, genes for T3SSYE1 of the predicted island Is.6 responded mainly to *in vitro* conditions. The differing transcriptional responses by the MH96 T3SSs depending on growth under *in vivo* and *in vitro* conditions allude to a specific role of each T3SS in MH96 pathogenicity. It was also noted that MH96 genes encoding the predicted PirAB binary toxin, RTX toxin and adenylate cyclase also responded to growth in culture, with significantly higher expression occurring *in vitro* compared *in vivo* conditions. Clearly some yet unknown environmental factor appears to control expression of T3SSYE1 and other insect active genes. It is plausible that abiotic factors like pH and concentration of oxygen or nutrients occurring during *in vitro* growth could mimic signals that MH96 is likely to encounter in its natural environment, such as intracellular replication within *G. mellonella* hemocytes (Hurst et al., 2015; Paulson, A.R., personal observation), infection of insect midgut or different insect hosts (Hurst et al., 2019, 2011) or even the occupation of the human urinary tract during asymptomatic catheter infection (Le Guern et al., 2018).

Along with T3SSs, another major class of well-studied VFs known from pathogenic *Yersinia* are adhesins, including T5SS-secreted adhesins and fimbriae (Chauhan et al., 2016). Putative adhesin genes upregulated by MH96 throughout or during specific stages of intrahemocoelic infection included T5SS-secreted AidA-like and filamentous hemagglutinin, and four-gene usher chaperone fimbrial cluster, the latter representing some of the most responsive genes to *in vivo* conditions in the entire MH96 genome. In addition to host cell attachment or biofilm formation, in other pathogenic *Yersinia* species, adhesins have also been linked to effector translocation by through enhanced attachment to host cells (Deuschle et al., 2015; Keller et al., 2015; Maldonado-arocho et al., 2013; Mühlenkamp et al., 2015). Perhaps adhesins also contribute to the effective anti-hemocytic activities of MH96 during intrahemocoelic infection of *G. mellonella* by too enhancing effector translocation efficiency like other pathogenic *Yersinia*, yet such mechanisms remain unknown among entomopathogenic bacteria.

The MH96 *in vivo* transcriptome revealed a high degree of functional redundancy, with multiple toxins, T3SS and T6SS effectors and adhesins all likely to target hemocytes during early and/or throughout multiple stages of infection. Such functional redundancy has also been observed in another highly entomopathogenic bacterium, *P. luminescens* (Wilkinson et al., 2009), that like MH96, produces many toxins, including insecticidal TC (ffrench-Constant et al., 2007), makes caterpillar floppy (Daborn et al., 2002), PirAB binary toxin (Waterfield et al., 2005; Wu & Yi, 2016) and several RTX-like toxins (Tobias et al., 2017). Such abundance of toxins carried within the genomes of obligate entomopathogenic bacteria is thought to have arisen by adaptation for over-kill of a wide variety of hosts (ffrench-Constant et al., 2003). This is likely also true for MH96, which has broad insecticidal activities against diverse insect hosts (Hurst et al., 2015, 2019, 2011, 2014). Also, MH96 is highly virulent against *G. mellonella* during intrahemocoelic infection but the LD_50_ of a Yen-TC deficient strain (ΔTC) was comparable to that of wild-type (∼ 3 cells). It was also found that both wild-type and ΔTC had the same ability to reduce the phenoloxidase activity of *G. mellonella* hemolymph 24 HPI, findings that implicate the presence of additional hemocytic VFs during intrahemocoelic infection by MH96 (Hurst et al., 2015). Transcriptional response by MH96 to *in vivo* conditions, including wide-spread and significant host-dependent upregulation of genes for Yen-TC components and numerous other predicted toxins, secretion systems, effectors and adhesins is another indicator that the highly insecticidal nature of MH96 stems from multiple diverse modes of action.

Beyond primary VFs that can act directly to modulate or destroy host cells, functional enrichment of the MH96 *in vivo* transcriptome also revealed changes in the temporal expression of genes for other putative in-host fitness factors (i.e., motility, metabolism, iron acquisition, response to host-induced stress, etc.) versus toxins, effectors and adhesins from early to late infection. Notably during the late stage of infection, MH96 significantly increased expression of genes related to flagellum and iron acquisition, compared to early and middle infection stages, which supports a role for iron acquisition and flagellum-related motility in MH96 in-host fitness during pre-cadaveric stages of insect infection.

Specific to entomopathogenic bacteria, both the flagellum and iron acquisition systems have already been implicated in intrahemocoelic infection of *G. mellonella* by triphasic nematode symbionts *P. luminescens* and closely-related *Xenorhabdus nematophila*. Flagellum-driven motility provided a selective fitness advantage during colonization of *G. mellonella* by *P. luminescens* (Easom & Clarke, 2008). Also, iron sensing by *X. nematophila* was shown to regulate the expression of flagellin gene *fliC* and hemolysin genes *xaxAB*, which significantly increased during late stage intrahemocoelic infection of *G. mellonella* and reached maximum levels following host death (Jubelin et al., 2011). Beyond entomopathogenic bacteria, iron homeostasis and hemoglobin utilization genes of *C. albicans* were found to be upregulated during later infection of *G. mellonella* and murine models compared to an earlier infection; however, in this study the early to late increase in iron-related genes was much greater in *G. mellonella* compared to the mouse model (Amorim-Vaz et al., 2015). Taken together, pathogens infecting *G. mellonella*, whether bacterial or fungal, appear to shift into an iron scavenging mode during late stage (i.e., pre-cadaveric) infection, indicating that pathogen sensing and response to *in vivo* iron fluxes during insect infection are likely an important component of in-host fitness or represent an interspecific competitive strategy.

## Acknowledgements

Research associate, Mitchell Weston, performed sample collection and RNA extraction for one of the *in vitro* samples used in this analysis. We would like to acknowledge Trystan Whang and Elena Kim for coordinating library preparation, quality control and sequencing at Macrogen Oceana. Finally, AgResearch Ltd. bioinformaticians Dr. Aurelie Laugraud, Dr. Charles Hefer and Paul Maclean helped with job execution on high-performance computing clusters.

## Author Contributions

Conceptualization, A.R.P., M.O., X.-X.Z., P.B.R., M.R.H.; Methodology, A.R.P., M.O., X.-X.Z., P.B.R., M.R.H.; Formal Analysis, A.R.P.; Writing – Original Draft Preparation, A.R.P.; Writing – Review & Editing, A.R.P., M.O., X.-X.Z., P.B.R., M.R.H.; Visualization, A.R.P.; Supervision, M.O., X.-X.Z., P.B.R., M.R.H.; Funding Acquisition, M.O., X.-X.Z., P.B.R., M.R.H. All authors have read and agreed to the pre-print version of the manuscript.

## Funding

This research aligned to and was supported by the Next Generation Biopesticides Programme, funded by the New Zealand Ministry for Business, Innovation and Employment (Contract C10×1310). Canadian National Science and Engineering Council for Postgraduate Scholarship-Doctoral Program and Universities New Zealand – Te Pōkai Tara via Massey University provided support in the form of student scholarships.

## Conflicts of Interest

The authors declare no conflicts of interest.

## Supplemental

**Table S1:**
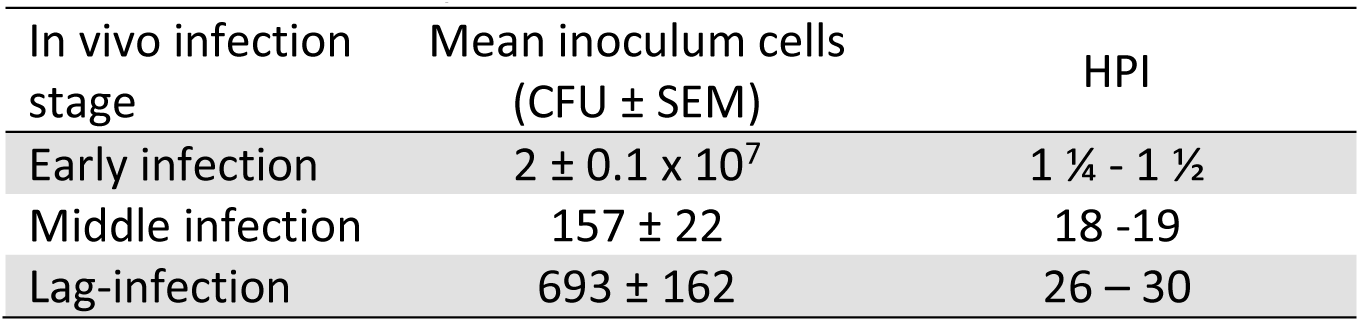
Mean number of *Yersinia entomophaga* MH96 cells from inoculum and hours-post infection (HPI) of *Galleria mellonella* by intrahemocoelic injection corresponding to lag, exponential and stationary in vivo growth phases at 25 °C. (n = 3 replicate plates). CFU = colony forming units, SEM = standard error of the mean and HPI = hours post infection.

| In vivo infection stage | Mean inoculum cells (CFU ± SEM) | HPI |
| --- | --- | --- |
| Early infection | $2 \pm 0.1 \times 10^7$ | 1 ¼ - 1 ½ |
| Middle infection | 157 ± 22 | 18 -19 |
| Lag-infection | 693 ± 162 | 26 – 30 |

**Table S2:**
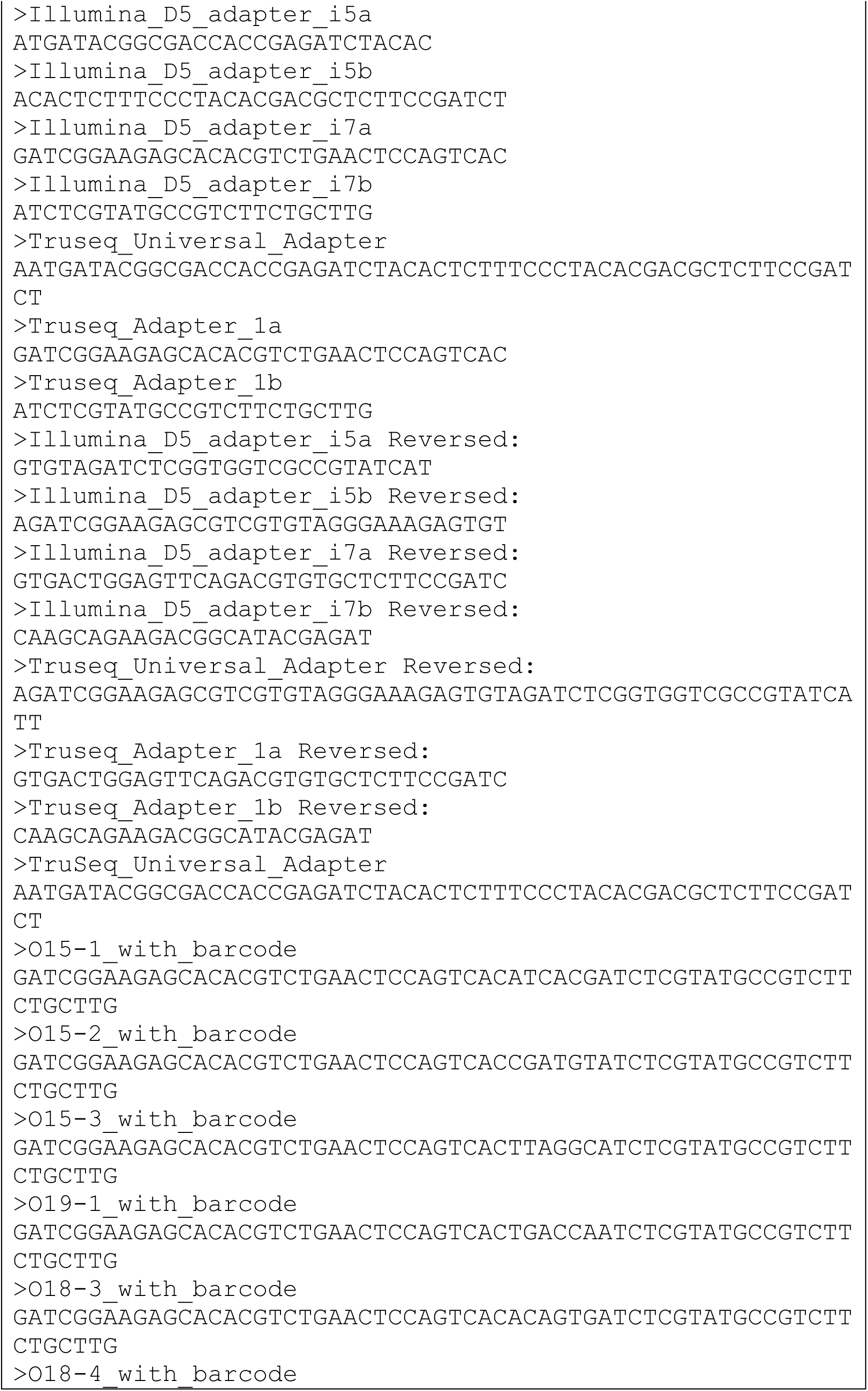
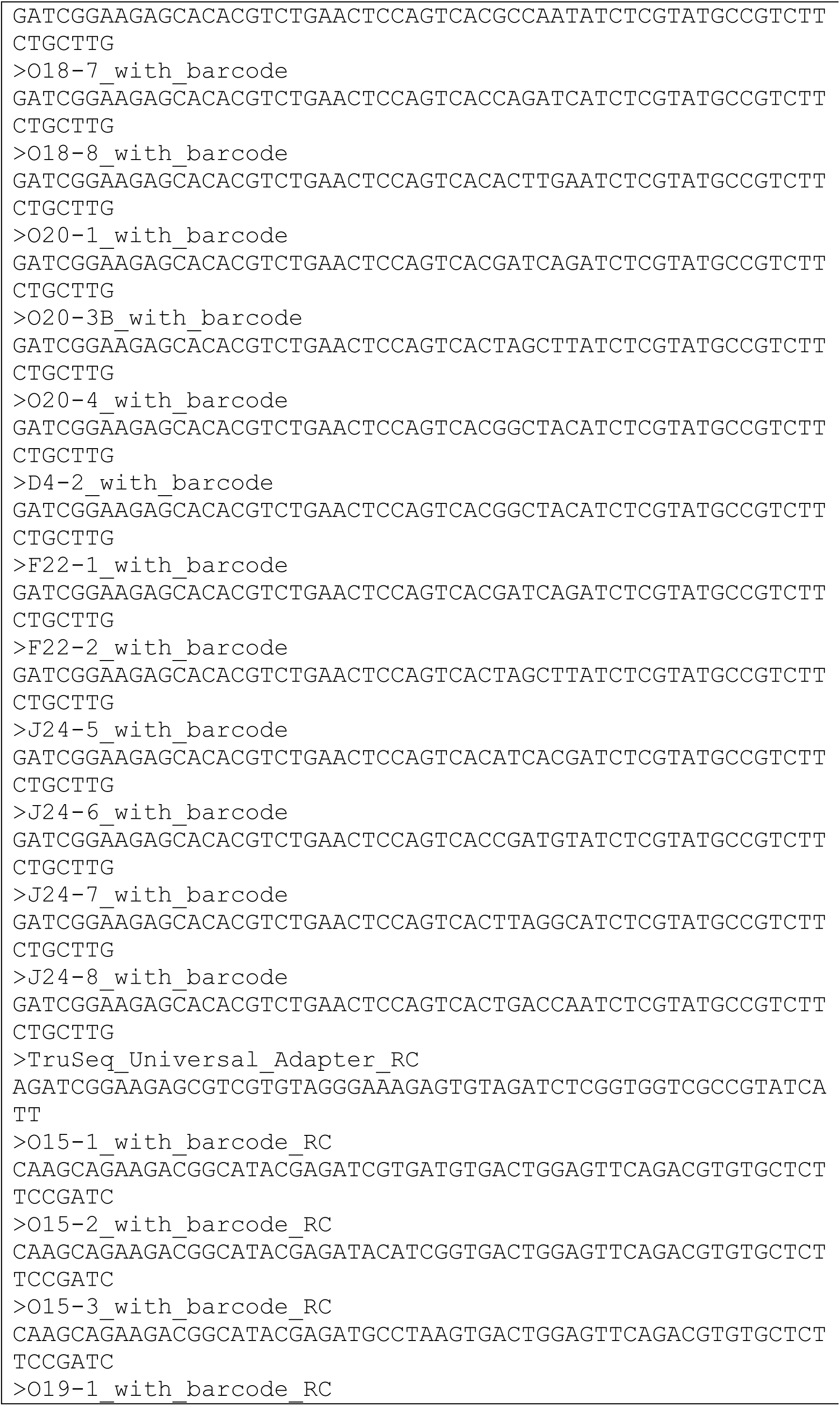
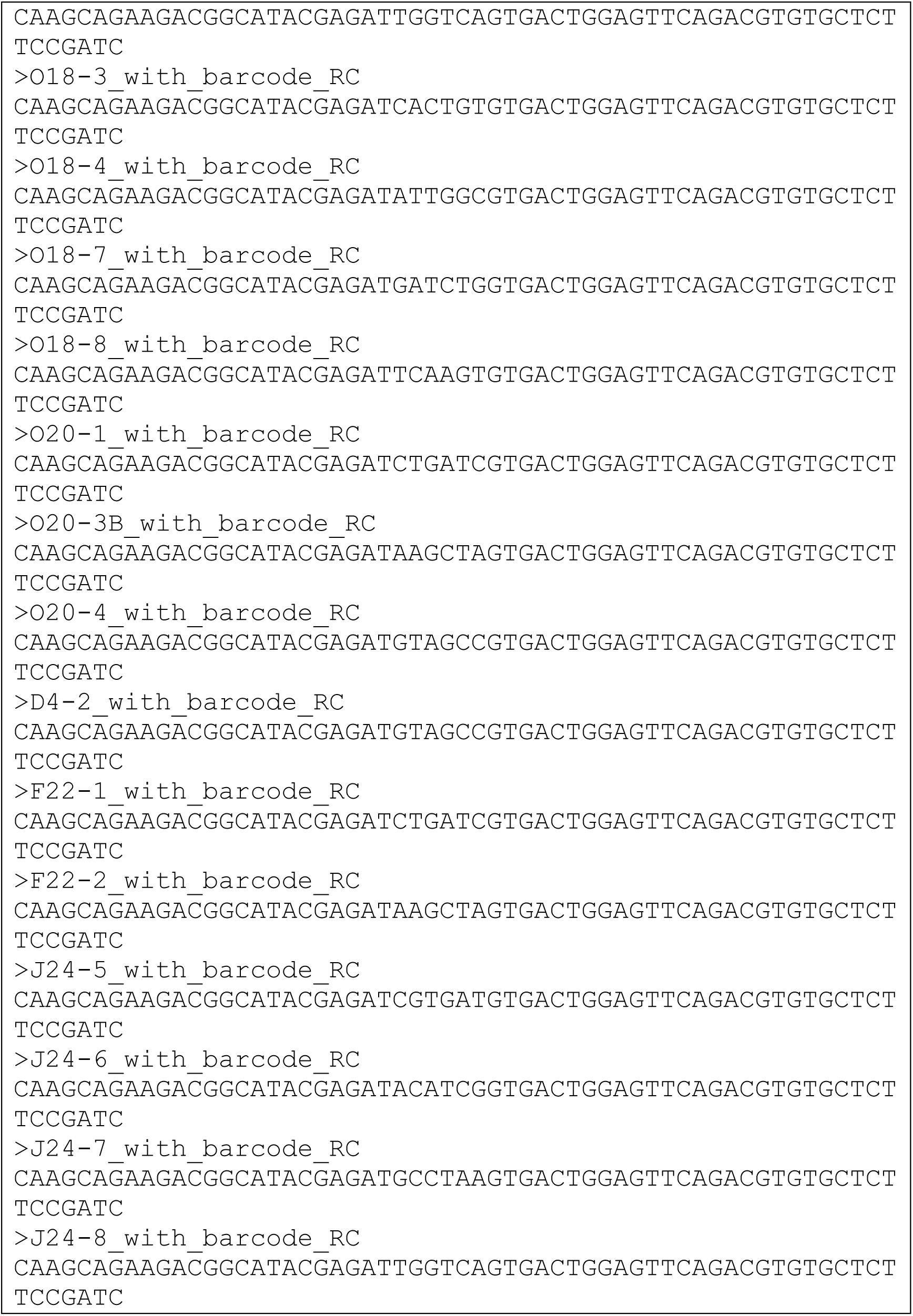
Adapter and barcode sequences used to trim all *in vivo* samples and *in vitro* samples ‘F22-1’ and ‘F22-2’ (exponential growth phase).

**Table S3:**
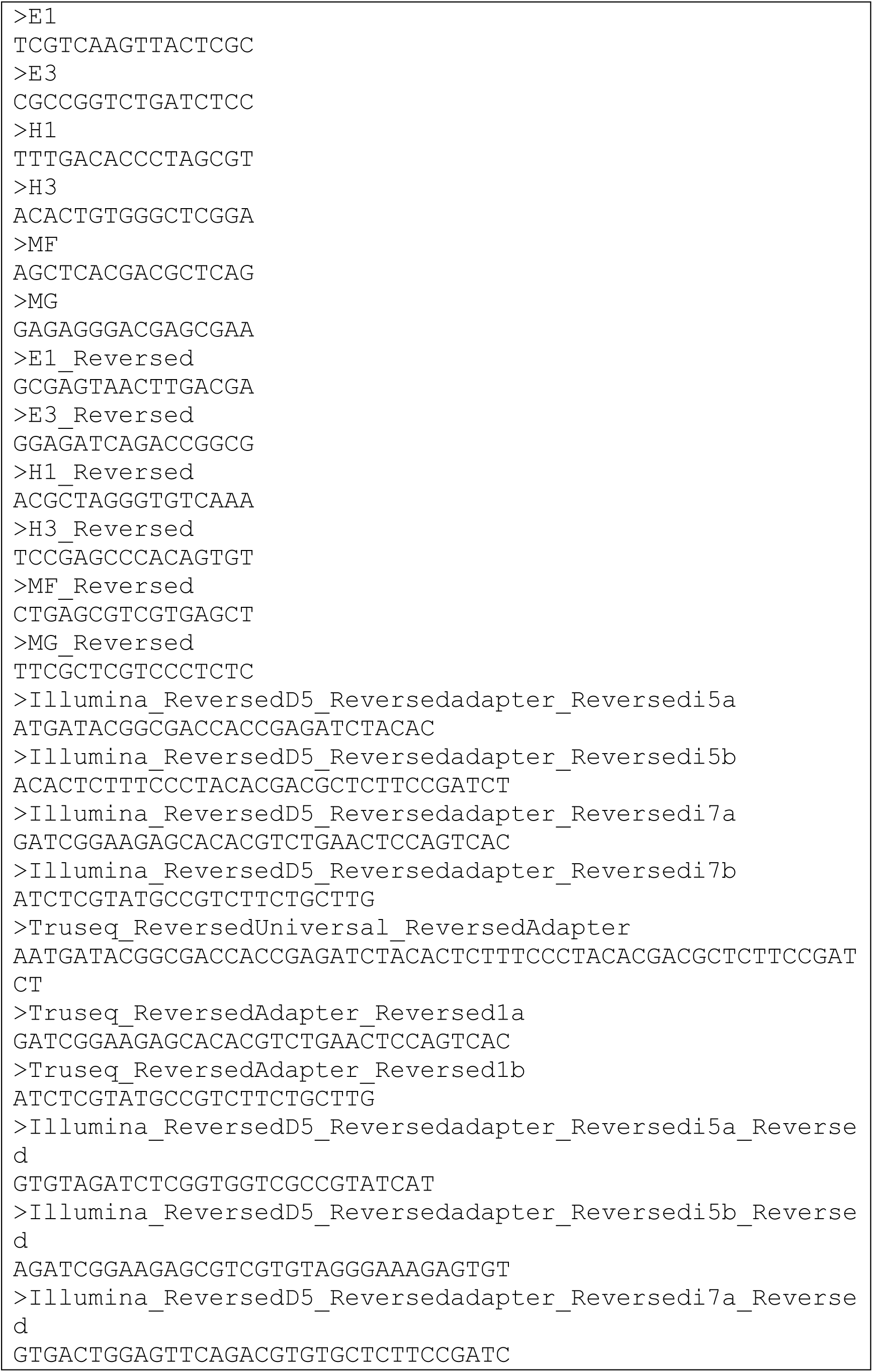
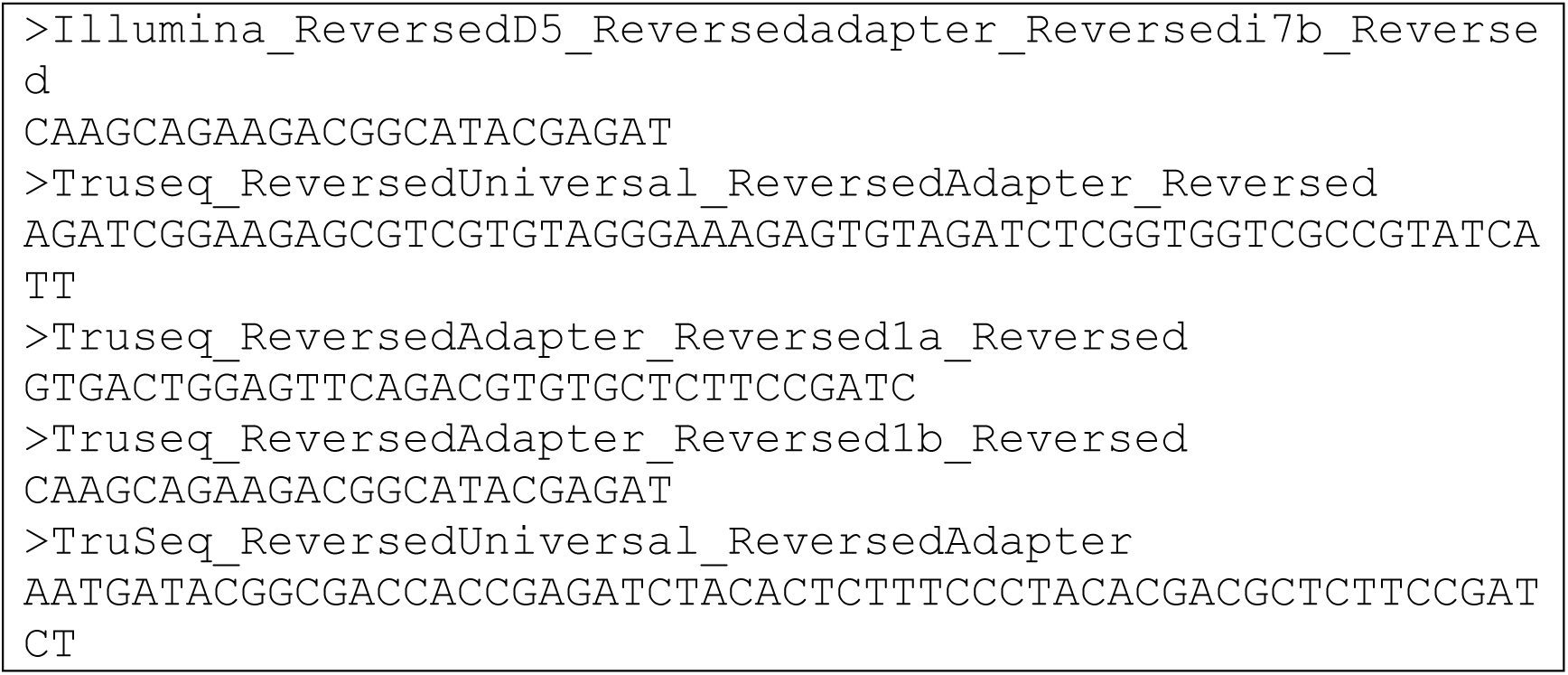
Adapter and barcode sequences used to trim all *in vitro* samples, except ‘F22-1’ and ‘F22-2’ (see Table S1).

**Table S4:** Categories of virulence factors considered in this study.

| Category | Definition | Examples |
| --- | --- | --- |
| *Primary – secreted and directly act against host cells/tissues* |  |  |
| Toxin | Proteins that are secreted and directly target host cells/tissues, giving rise to pathogenesis. | Insecticidal toxin complex, vegetative insecticidal protein or Cry toxin. |
| Effector | Proteins that are translocated directly into host cells by type 3 or 6 secretion systems. | *Yersinia* spp. Yop or *Salmonella* spp. Spa type 3 secretion system and hemolysin co-regulated protein (Hcp) or polymorphic rearrangement hot-spot (Rhs) toxins. |
| *Secondary – all other non-primary factors contributing to pathogenesis (i.e., gene abrogation results in attenuated virulence).* |  |  |
| Adherence | External structures used to attach to host cells or tissues. | Fimbriae or filamentous hemagglutinins. |
| Defense response | Factors used to detoxify against antimicrobial compounds or host-induced stress. | Superoxide dismutase, efflux pumps, iron-sulfur cluster. |
| Horizontal gene transfer | Factors that promote lateral gene acquisitions or mobilization. | Conjugative pili or integrases. |
| Iron acquisition | Systems used by pathogenic bacteria to acquire iron in the host environment. | Siderophores, hemolysins and iron-specific transporters. |
| Metabolic adaptation | Enzymes and transporters involved in bioconversion of host-derived nutrients. | Amino acid transporters or chitinases. |
| Mobility | Factors related to motility. | Flagella or chemotaxis. |
| Outer membrane | Factors related to modification of outer membrane/capsule often used to evade host-immune response. | Lipopolysaccharide or outer membrane proteins. |
| Regulation | Regulatory factors controlling pathogenic response to host. | Two-component sensors, quorum sensing or diguanylate cyclase. |
| Secretion system | Multi-component systems used for translocation and secretion of toxins and effectors by bacteria. | Type 2, 3, and 6 secretion systems. |
| Unclassified | Factors that could not be classified specific categories listed above. | Non-ribosomal peptide synthetase or tellurium resistance genes. |

Table S5. Differentially expressed genes identified from *Yersinia entomophaga* MH96 for time-resolved *vivo* vs. *in vitro* comparison from upper quartile normalized count data using *Limma* with Benjamini and Yekutieli adjusted p-value cutoff of 0.05.

[See “Supplemental Table S5 - differentially expressed genes identified from Yersinia entomophaga time-resolved infection-series.xls” in the supplemental zip folder].

Table S6. In-host putative virulence factors identified in *Yersinia entomophaga* MH96 using fuzzy clustering and virulence annotations from VFDB.

[See “Supplemental Table S6 - In-host putative virulence factors identified in Yersinia entomophaga using FCM and VFDB.xls” in the supplemental zip folder].

**Figure S1:**
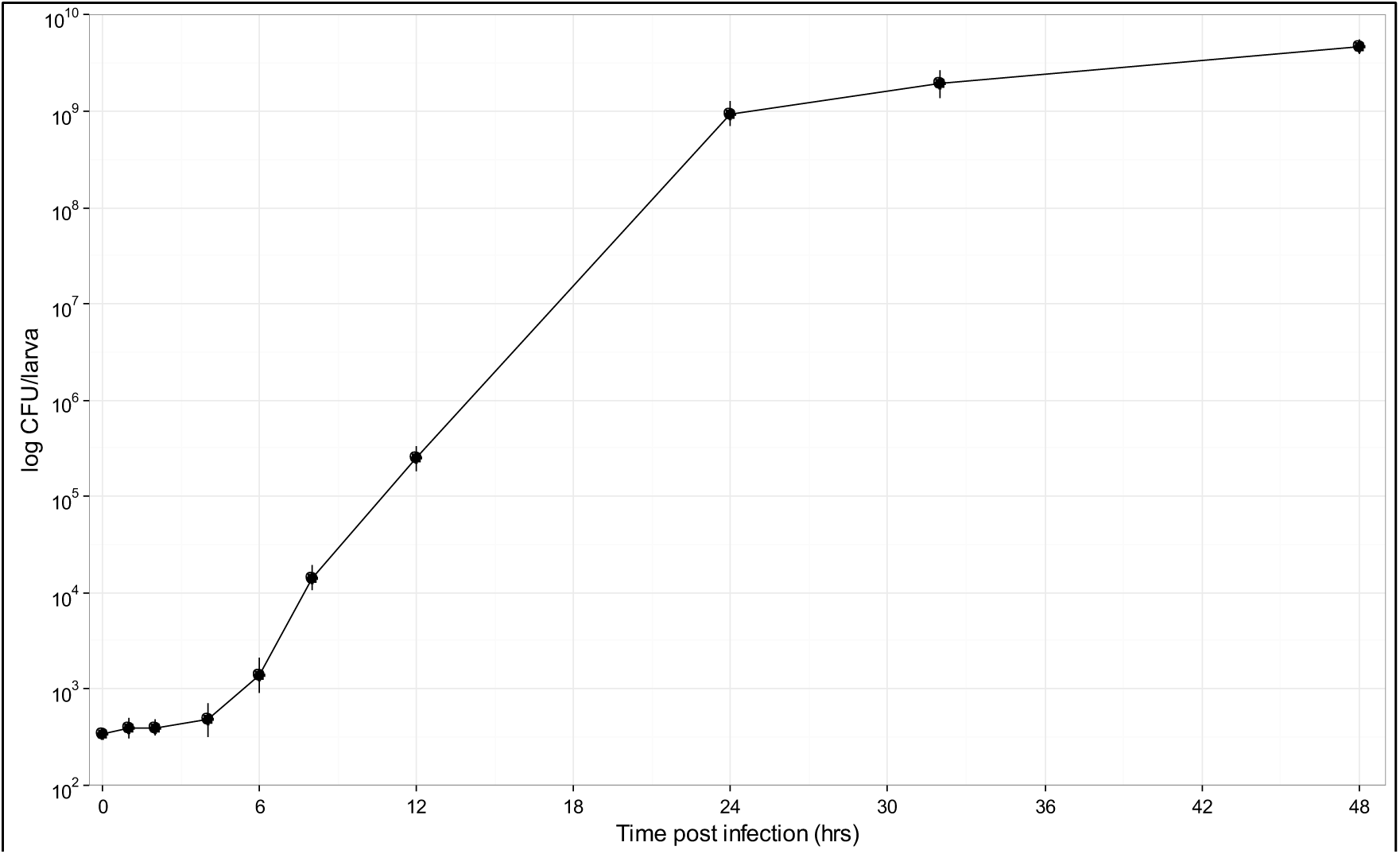
*In vivo* growth curve of *Yersinia entomophaga* MH96 in the hemolymph of *Galleria mellonella* at 25 °C. Larvae were injected with approximately 345 cells in 10 μl PBS (time-point 0) and CFU/larva were enumerated from 3 larvae per timepoint at 1, 2 4, 6, 8, 12, 24, 32 and 48 hours post-infection. Points represent mean density and vertical bars represent standard error.

**Figure S2.**
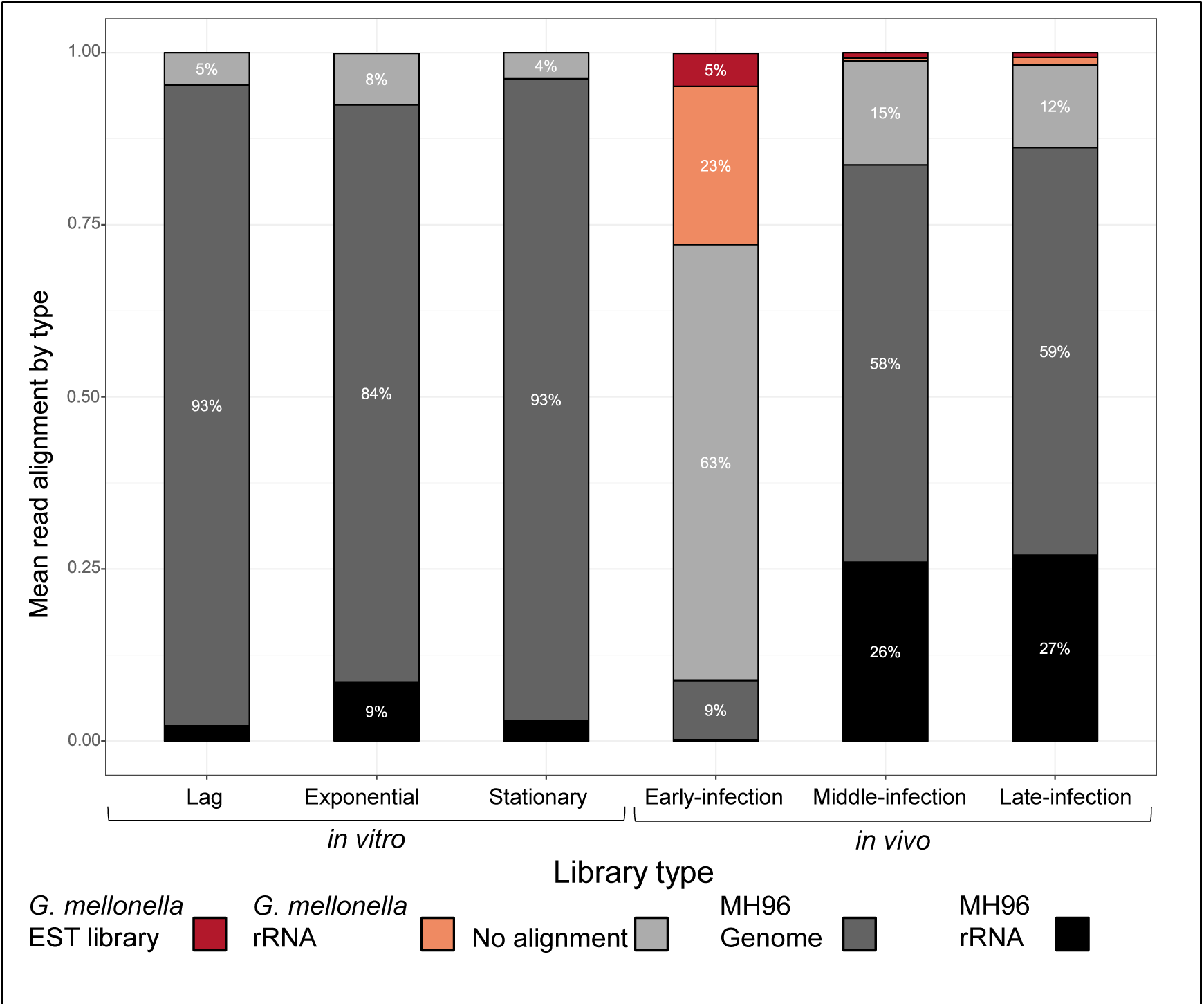
Mean proportion of paired-end reads attributed to host and pathogen based on average percent alignment to *Galleria mellonella* rRNA and EST sequence or *Yersinia entomophaga* MH96 genome or rRNA sequence (*in vivo:* n = 4, *in vitro*: n = 2). The growth phase and estimated cell number (CFU/g and CFU/ml for *in vivo* and *in vitro*, respectively) are provided in Table 1). Only percentages of 4 % and greater are shown.

**Table S7:** Mean percentage of trimmed paired-end reads aligning to *Yersinia entomophaga* MH96 reference genome, including protein-coding, tRNA and non-coding regions (*in vivo:* n = 4, *in vitro*: n = 2).

| Infection-stage / growth-phase | Total align-ments | Protein coding region | Anti-sense protein coding region | tRNA | Non-annotated regions |
| --- | --- | --- | --- | --- | --- |
| *In vivo* |  |  |  |  |  |
| Early infection | 9 % | 77 % | 2 % | 1 % | 22 % |
| Middle infection | 58 % | 74 % | 2 % | 1 % | 24 % |
| Late infection | 59 % | 75 % | 1 % | 0 % | 24 % |
| *In vitro* |  |  |  |  |  |
| Lag | 93 % | 84 % | 2 % | 1 % | 14 % |
| Exponential | 84 % | 83 % | 1 % | 1 % | 15 % |
| Stationary | 93 % | 85 % | 1 % | 0 % | 14 % |

**Figure S3:**
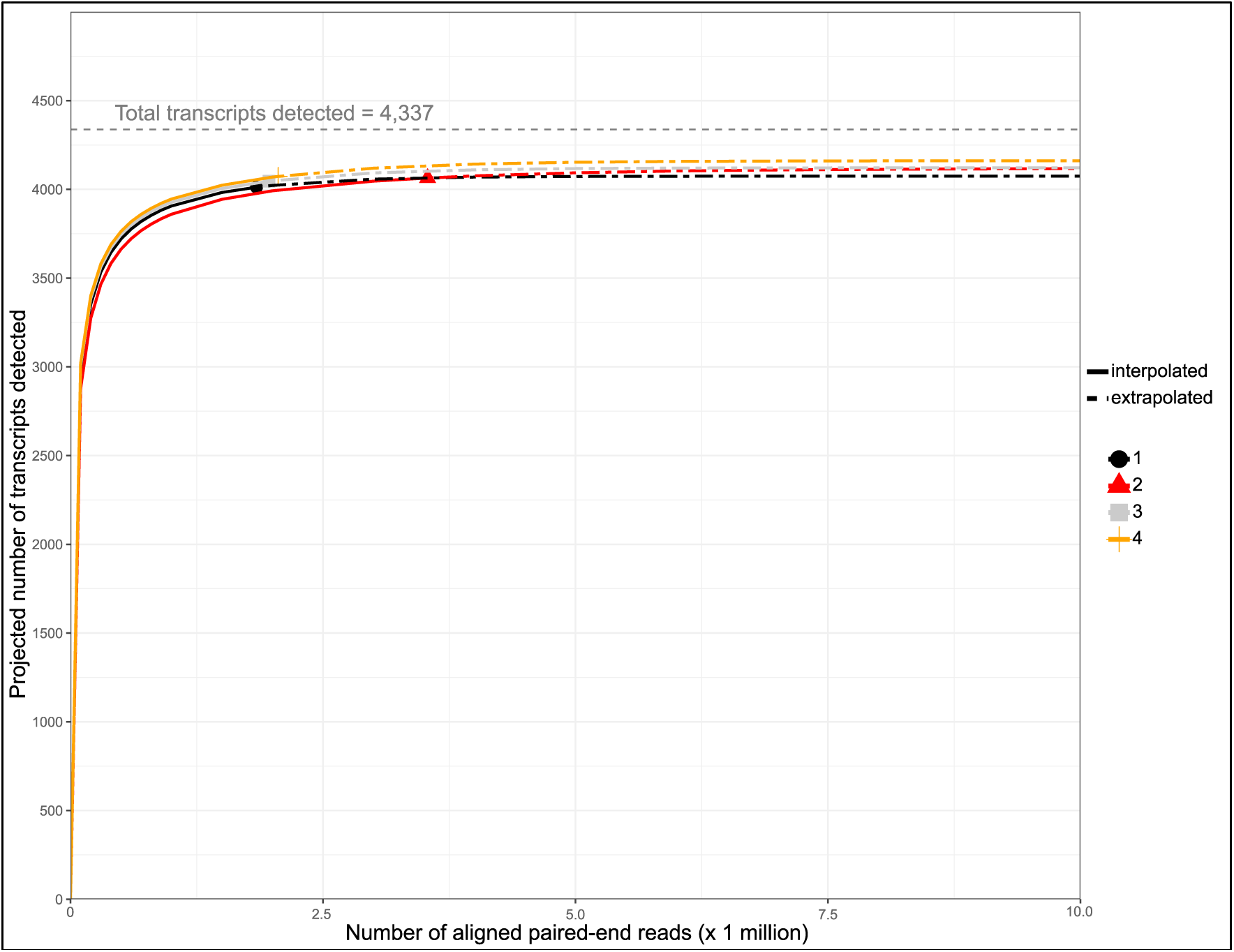
Predicted transcript detection based on the number of aligned paired-end reads for *in vivo* early infection libraries by rarefaction analysis. 1, 2, 3 and 4 represent individual libraries.

**Figure S4.**
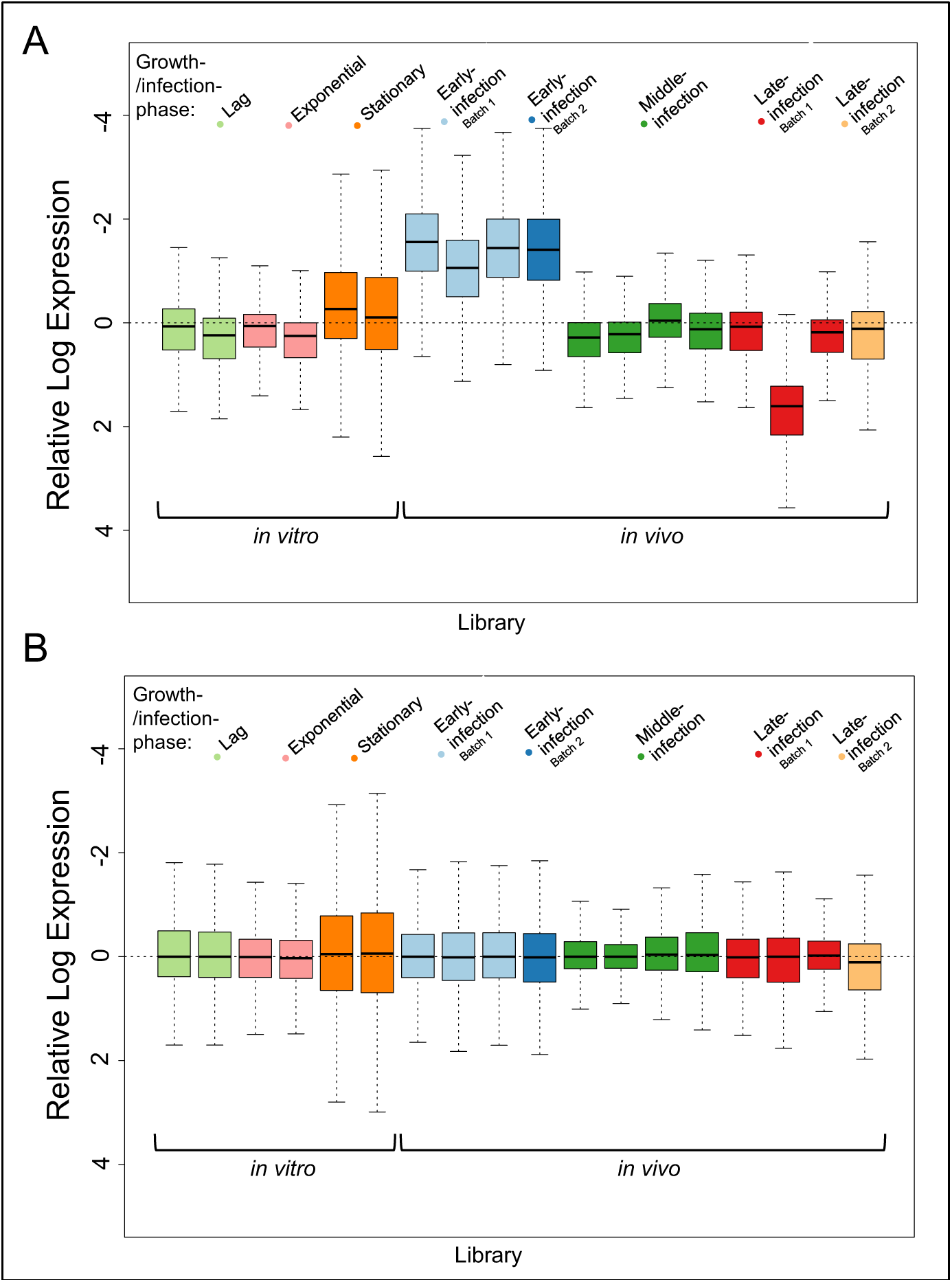
Relative log expression of *in vivo* and *in vitro* libraries represented as median boxplots. A) unnormalized and B) upper-quartile normalized. Individual collection batches are noted where applicable.

**Figure S5.**
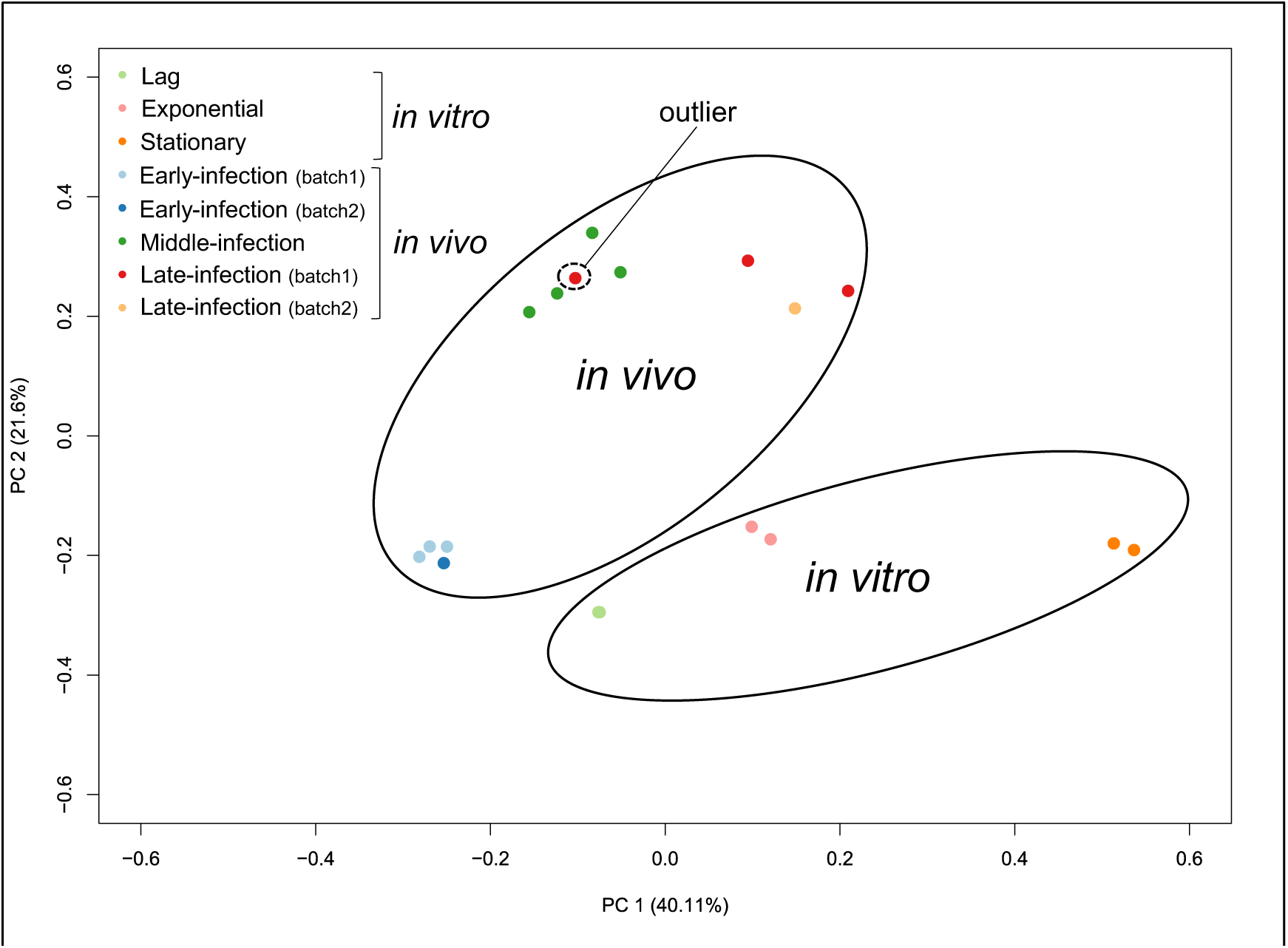
Principle component analysis of *in vivo* and *in vitro* library upper quartile normalized count data. Outlier late infection *in vivo* sample is indicated and was removed from subsequent analysis.

**Figure S6.**
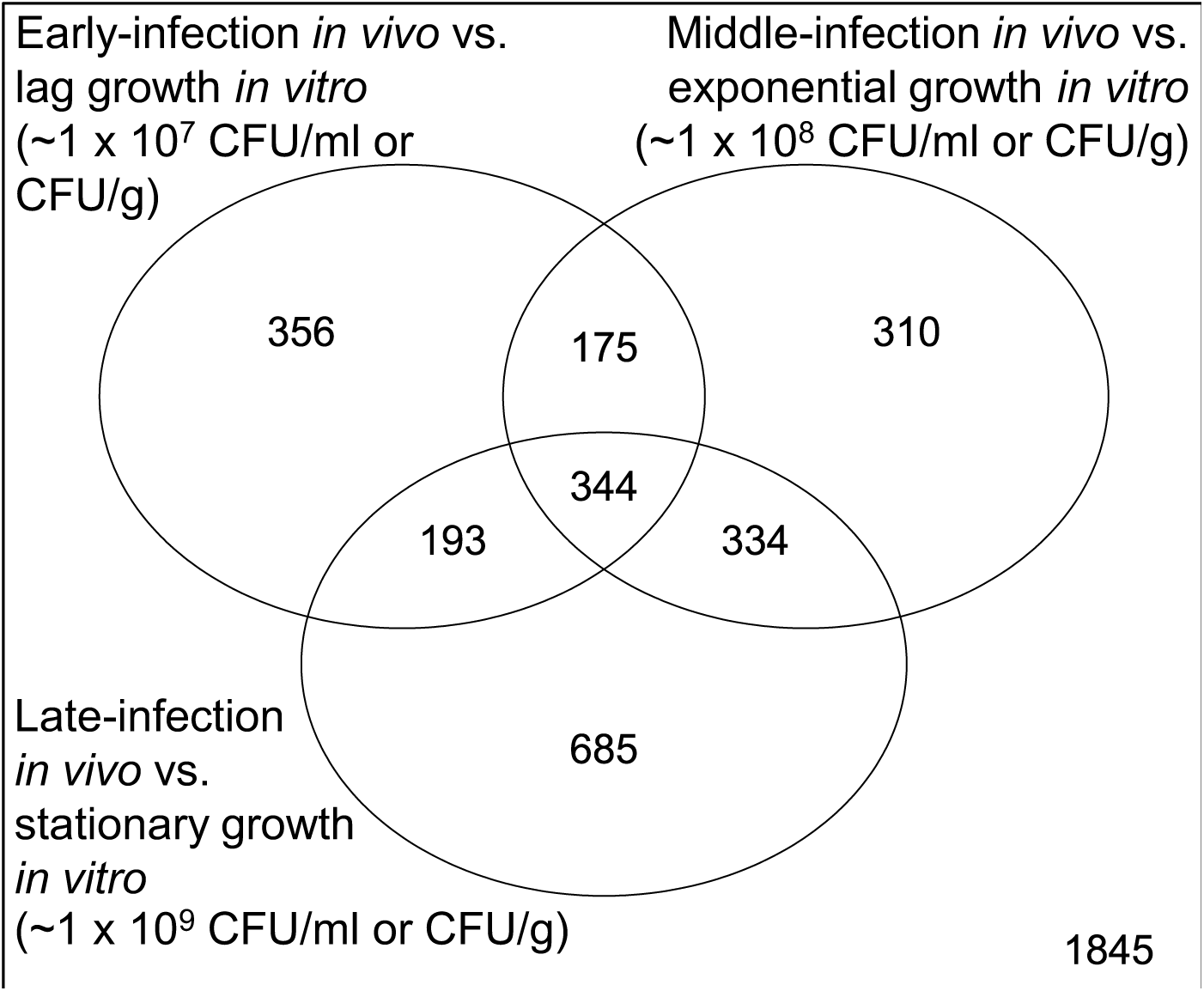
Differentially expressed transcripts from *in vivo* or *in vitro* libraries during early, middle and late infection and lag, exponential and stationary growth phases, respectively. Significance was multiple-test corrected (Benjamini & Yekutieli, 2001) with an adjusted p-value threshold of 0.05. See supplemental Table S5 for more details on the contrasts represented in the Venn groups.

**Table S8:**
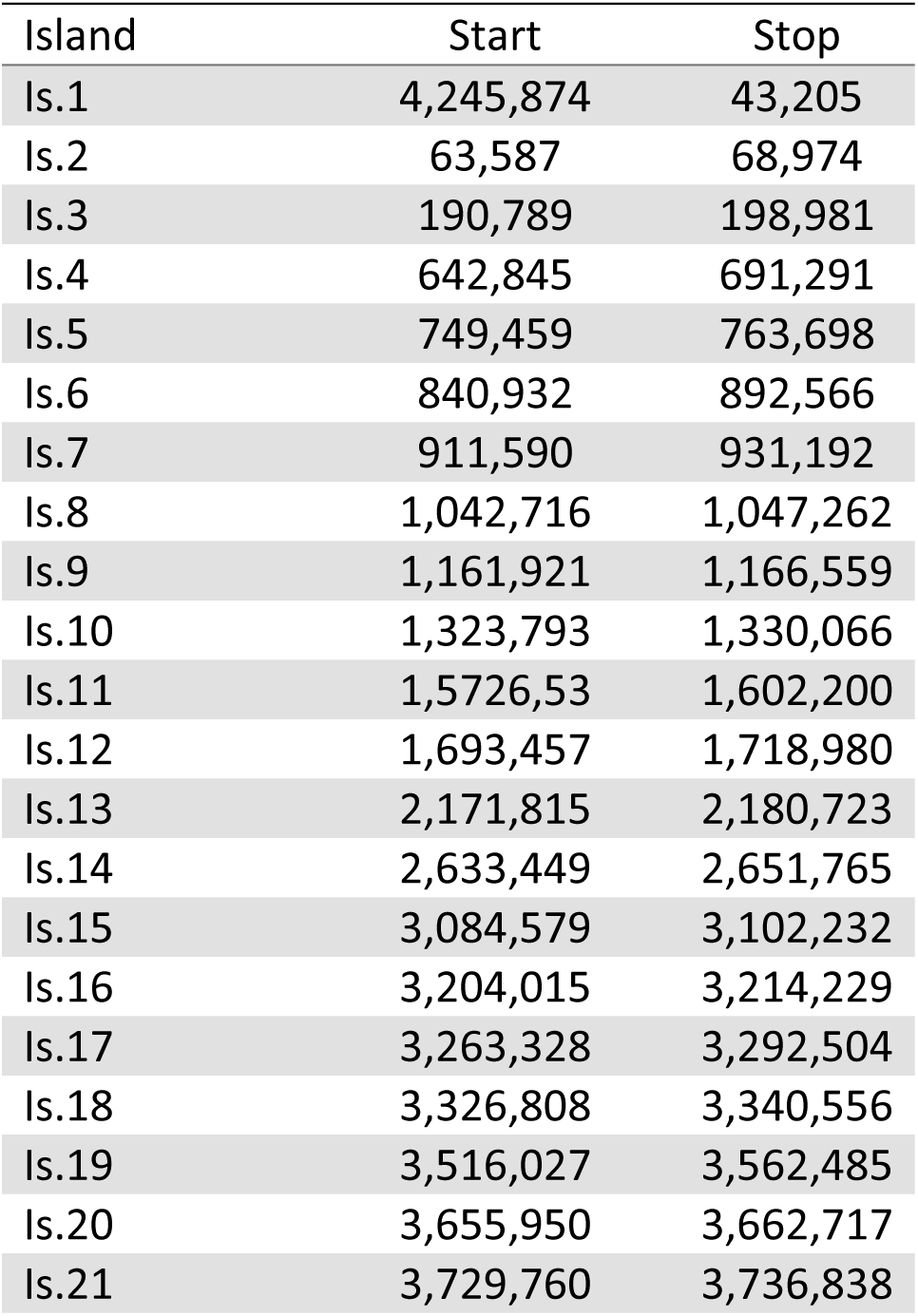
Location of predicted genomic islands in the genome of *Y. entomophaga* MH96 based on Island Viewer 4 analysis.

**Figure S7:**
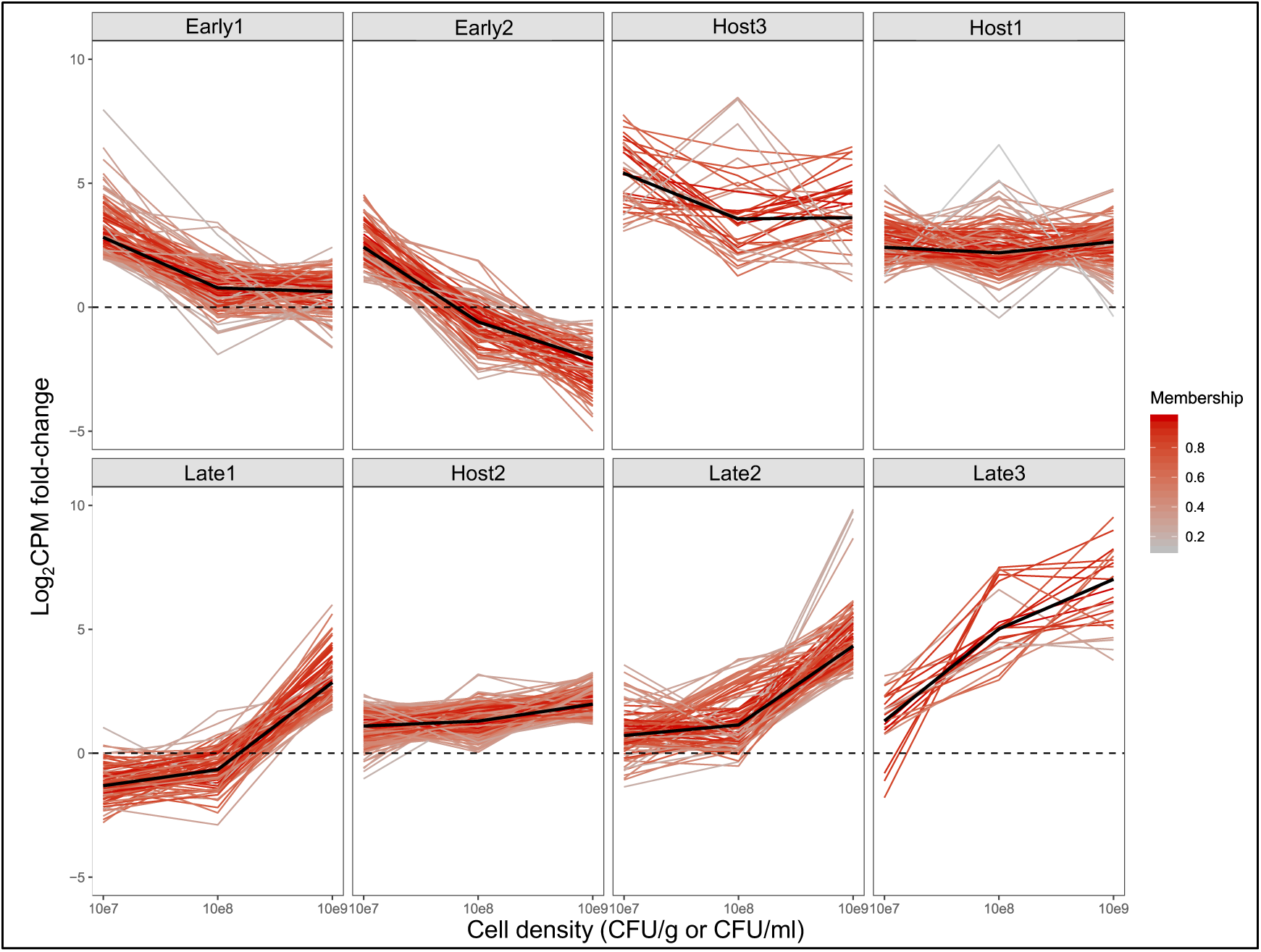
Host-specific transcripts from *Yersinia entomophaga* MH96 *in vivo* RNA-seq identified using c-means fuzzy clustering algorithm (k = 20, m = 1.5). Log_2_ counts-per-million (CPM) fold-change between *in vivo* and *in vitro* samples for each cell density. Black line = median log_2_CPM fold-change. CFU = colony forming unit.

## Notes

https://www.ncbi.nlm.nih.gov/geo/query/acc.cgi?acc=GSE142509

https://github.com/damselflywingz/In_vivo_infection_series_fuzzy_RNA-seq

### Competing Interest Statement

The authors have declared no competing interest.

